# Spatial regulation of clathrin-mediated endocytosis through position-dependent site maturation

**DOI:** 10.1101/834523

**Authors:** Ross TA Pedersen, Julian E Hassinger, Paul Marchando, David G Drubin

## Abstract

During clathrin-mediated endocytosis (CME), over 50 different proteins assemble on the plasma membrane to reshape it into a cargo-laden vesicle. It has long been assumed that cargo triggers local CME site assembly in *Saccharomyces cerevisiae* based on the discovery that cortical actin patches clustered near exocytic sites are CME sites. Quantitative imaging data reported here lead to a radically different view of which CME steps are regulated and which steps are deterministic. We quantitatively and spatially describe progression through the CME pathway and pinpoint a cargo-sensitive regulatory transition point that governs progression from the initiation phase of CME to the internalization phase. Thus, site maturation, rather than site initiation, accounts for the previously observed polarized distribution of actin patches in this organism. While previous studies suggested that cargo ensures its own internalization by regulating either CME initiation rates or frequency of abortive events, our data instead identify maturation through a checkpoint in the pathway as the cargo-sensitive step.

**Summary:** Pedersen, Hassinger, et al. investigate steps of the clathrin-mediated endocytosis pathway that are subject to regulation. They report position-dependent differences in endocytic site maturation rates in polarized cells and suggest that cargo controls endocytic internalization through tuning site maturation rather than site initiation.

## Introduction

Membrane trafficking processes require coordinated cargo capture and membrane reshaping. One such process is clathrin-mediated endocytosis (CME), which is the major trafficking route into the cell (Bitsikas et al., 2014). Numerous studies have suggested that cargo influences the rate of CME by modulating the proportion of endocytic sites that continue to completion as opposed to aborting precociously (Loerke et al., 2009; Mettlen et al., 2010; Liu et al., 2010; Ehrlich et al., 2004). However, this proposed mode of coordinating cargo capture with endocytosis was called into question when studies of genome-edited mammalian cells did not reveal widespread abortive CME events (Doyon et al., 2011; Hong et al., 2015). In the budding yeast *Saccharomyces cerevisiae*, it is widely accepted that the vast majority of CME events proceed to completion (Kaksonen et al., 2003, 2005), but regulation of CME progression is nevertheless apparent. The observation in budding yeast that internalizing CME sites (originally observed as “actin patches”) are clustered near sites of polarized exocytosis led the field to assume that cargo delivered by exocytosis locally promotes CME site initiation, but cargo-dependent acceleration of CME initiation has not been directly observed. (Godlee and Kaksonen, 2013; Goode et al., 2015; Field and Schekman, 1980; Adams and Pringle, 1984; Kaksonen et al., 2003). Thus, how the CME pathway is regulated, and which steps are subject to regulation, remain unresolved.

*S. cerevisiae* is an ideal organism for studies of the CME pathway. Aside from practical advantages such as simplicity, tractable genetics, and accessible genome editing*, S. cerevisiae* has a remarkably regular CME pathway, and many of the molecular components of the CME machinery are conserved from budding yeast to humans (Kaksonen et al., 2003, 2005, 2006). Over 50 proteins are recruited to every CME site in budding yeast with an invariant order and with fairly regular kinetics (Lu et al., 2016). This highly stereotyped CME pathway in *S. cerevisiae* is ideal for determining which CME steps are deterministic and which are subject to regulation.

Budding yeast CME proteins can be organized conceptually into functional modules based on shared timing of recruitment and dynamic behavior, common functions, and spatial organization at CME sites (Fig. 1A, Carroll et al., 2012; Kaksonen et al., 2005; Lu et al., 2016). Briefly, proteins of the early and early coat modules initiate CME sites (Lu and Drubin, 2017; Stimpson et al., 2009; Godlee and Kaksonen, 2013). Next, intermediate coat proteins, which bind to actin filaments and couple actin growth to membrane invagination, are recruited (Skruzny et al., 2012; Carroll et al., 2012). Late coat proteins arrive and recruit actin assembly machinery consisting of proteins of the WASP/Myosin complex (Sun et al., 2015; Bradford et al., 2015). The WASP/myosin complex orchestrates a critical burst of actin assembly and myosin activity to bend the plasma membrane into a pit (Lewellyn et al., 2015; Pedersen and Drubin, 2019; Sun et al., 2006; Manenschijn et al., 2019). While the early coat, intermediate coat, and late coat proteins internalize with the plasma membrane, the early module proteins dissociate from CME sites at the time of actin assembly (Carroll et al., 2012). Actin filaments recruit a host of actin binding proteins and disassembly factors collectively referred to as the actin module. At the same time, proteins of the scission module arrive to release the nascent vesicle. The timing and modular organization of this process are conserved from yeast to mammals (Taylor et al., 2011).

**Figure 1:**
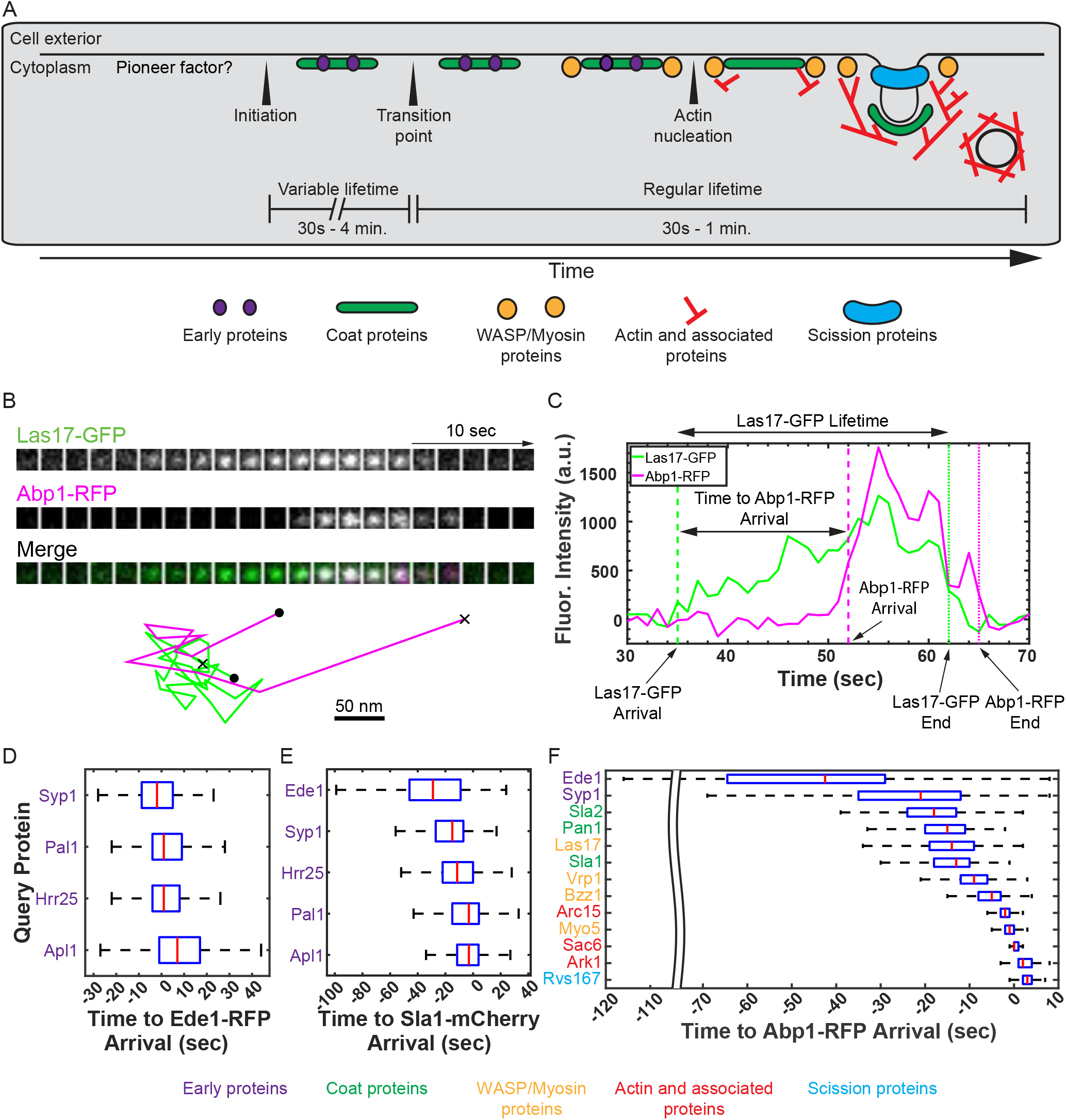
Quantitatively probing the budding yeast clathrin-mediated endocytosis pathway through systematic imaging. (A) Illustration of key steps in the budding yeast CME pathway. Mechanisms of site initiation, transition from the early, variable lifetime stages to the later, more regular lifetime stages, and initiation of actin assembly are poorly understood. Diagram adapted from Tonikian et al., 2009. (B) Top: montage of Las17-GFP and Abp1-RFP recruitment to an endocytic site viewed *en face* by live two-color TIRF microscopy of budding yeast. Bottom: centroid position tracks in the x and y dimensions for the Las17-GFP (green) and Abp1-RFP (magenta) spots depicted in the montage. • - beginning, x – end. See Video 1. (C) Fluorescence intensity vs. time for Las17-GFP and Abp1-RFP spots depicted in panel B. Timepoints of note and key measurements recorded are indicated by dotted lines and arrows, respectively. (D – F) Box and whisker plots of the time from the appearance of the indicated GFP-tagged query protein to the appearance of (D) Ede1-RFP, (E) Sla1-mCherry, and (F) Abp1-RFP. Red lines are median values, boxes indicate interquartile range, and whiskers indicate full range of the data. Colored text identifies each protein imaged as a component of the Early, Coat, WASP/Myosin, Actin, or Scission modules according to the color scheme below.

Despite this detailed knowledge of the CME machinery, how key transitions in the process are triggered remains poorly understood (Fig. 1A). The budding yeast early and early coat proteins persist at CME sites for a variable period of time before recruitment of the late arriving coat components, which are reported to have more regular lifetimes (Carroll et al., 2012; Stimpson et al., 2009; Peng et al., 2015; Newpher et al., 2005). A previous study provided preliminary evidence consistent with cargo regulating the transition from the “variable phase” to the “regular phase”, but this model was not thoroughly tested, the point in the pathway that is regulated was not identified, and the physiological implications of the model were not examined (Carroll et al., 2012).

Here, we performed systematic live-cell imaging analyses of proteins in the budding yeast CME pathway. Our data quantitatively separate the CME pathway into an early phase and a late phase. The behavior of proteins in each phase predicts the behavior of others in the same phase, but not the behavior of proteins in the other phase. Further experiments lead us to propose that the presence of cargo accelerates the transition from the early phase to the late phase, so this regulatory transition is likely a cargo checkpoint. We propose that endocytic sites are initiated widely but stall in the absence of cargo, maturing to the late, internalization phase more readily in the presence of cargo.

## Results

### Systematic live-cell imaging of clathrin-mediated endocytosis reveals inherent variability in timing and abundance of protein recruitment

We used systematic imaging to quantitatively describe progression through the CME pathway. While each of the proteins of the CME pathway we examined had been imaged previously to determine a recruitment order, we performed a deeper quantitative analysis to uncover previously undetected features and relationships among the proteins. Recruitment order and timing in the budding yeast CME pathway were previously determined by many pair-wise imaging iterations using two-color fluorescence microscopy. Here we imaged CME protein recruitment timing relative to three reference proteins, providing an orthogonal approach to determining the recruitment order and relative timing.

We acquired live-cell two-color total internal reflection fluorescence microscopy (TIRFM) movies of budding yeast strains expressing nearly two-dozen different fluorescent fusion proteins (Table S1). Each yeast strain encoded an endocytic query protein tagged with GFP and a reference protein for various stages of progression through CME tagged with a red fluorophore. All proteins were endogenously tagged and expressed from their own promoters in the genome to avoid overexpression artifacts. We chose the Eps15 homologue Ede1, a key component of the early endocytic module (Stimpson et al., 2009; Carroll et al., 2012; Lu and Drubin, 2017), as a reference protein to report on CME site initiation. Sla1, a late-arriving endocytic coat protein (Kaksonen et al., 2003), was used as a reference protein for the transition from the variable stage of CME to the regular stage. Finally, we chose the actin binding protein Abp1, a reporter of endocytic actin networks in budding yeast (Kaksonen et al., 2003), to report on initiation of actin assembly. All of these reference proteins are relatively abundant at endocytic sites (and hence bright). We chose TIRFM in lieu of the more standard imaging modality of widefield microscopy in a medial focal plane of yeast cells (Kaksonen et al., 2003, 2005). TIRFM has a superior signal-to-noise ratio as well as minimal photobleaching, although this comes at the cost of losing the ability to track the endocytic proteins once they move away from the plasma membrane during internalization. The benefits of TIRFM were critical for faithfully detecting and tracking early endocytic proteins, because they produce a dim, long-lived signal at CME sites. By imaging different GFP-tagged query proteins in two-color movies with common reference markers, we were able to compare the dynamics of a large subset of the CME proteins at endocytic sites.

To chronicle progression through the CME pathway, we chose GFP-tagged query proteins from each endocytic module. We imaged the early module proteins Syp1 (an FCho homologue) and Hrr25 (a casein kinase homologue) and the early coat proteins Pal1 and Apl1 (a component of the AP-2 complex) with reference to the early protein Ede1 to capture the earliest stages of CME. We also imaged the early module proteins Ede1, Syp1, and Hrr25 and the early coat proteins Pal1 and Apl1 with reference to the late coat protein Sla1 to monitor the transition from the early stage of CME to the late stage. Finally, we imaged a large sample of CME proteins with reference to Abp1 to monitor the late stage of CME and the initiation of actin assembly in particular. We used Ede1 and Syp1 as representative early proteins; Sla2 (human homologue: Hip1R) as a model intermediate coat protein; Pan1 (intersectin) and Sla1 as model late coat proteins; Las17 (Wiscott-aldrich syndrome protein, WASP), Bzz1 (toca1), Vrp1 (WASP interacting protein, WIP), and Myo5 (myosin 1e) as model WASP/Myosin module proteins; Arc15 (an Arp2/3 complex subunit), Sac6 (fimbrin), and Ark1 (Cyclin G-associated kinase/Adaptor-associated kinase) as model actin module proteins; and Rvs167 (amphiphysin) as a model scission module protein. Movies of these query proteins yielded new information about every stage of the CME pathway.

A combination of established and custom MATLAB scripts was used to monitor fluorescently tagged proteins at CME sites. CME sites are smaller than the diffraction limit of fluorescence microscopy, so they appear as ∼200 nm diameter spots in our movies. Fig. 1B depicts an example montage of an endocytic site. In this case, the GFP-tagged query protein Las17, arrives before the reference protein Abp1-RFP. Both proteins then disappear as the endocytic pit internalizes and leaves the TIRFM field (See also Video 1). We used the cmeAnalysis MATLAB package to automatically track all of the endocytic sites in many fields of yeast cells (Aguet et al., 2013). We then used custom MATLAB scripts to: identify and associate colocalized green and red spots, reject CME sites that were too close to one another to resolve independently, and extract background-subtracted fluorescence intensities for each fluorescence channel at each site (See Materials and Methods for full details). Using our software, we tracked individual endocytic sites from their initiation to their disappearance and extracted time-resolved fluorescence intensity data for each channel (Fig. 1B-C).

Our systematic imaging data set independently supports the previously proposed recruitment order for CME proteins while providing additional mechanistic insights. We generated plots of fluorescence intensity vs. time for each fluorescently-tagged protein at each tracked CME site and recorded lifetimes, maximum fluorescence intensities, and the delay between query protein arrival and reference protein arrival (Fig. 1C). When we ordered the proteins using our time to reference protein arrival data, we recapitulated the previously deduced recruitment order (Fig. 1D – F, Tables S2-4). First, our plots show that all early arriving proteins (Ede1, Syp1, Pal1, Hrr25, and Apl1) arrive at CME sites at about the same time when imaged with reference to Ede1-RFP (Fig. 1D, Carroll et al., 2012). When imaged with reference to Sla1-mCherry, Ede1 appears to be the earliest arriving protein, although this is likely at least in part due to Ede1 being the brightest of the group, which unavoidably results in earlier signal detection (Fig. 1E, also see Fig. S1A). Our dataset using Abp1 as a reference protein recapitulates the canonical timeline of the CME pathway, an ordering which was established through painstaking sequential rounds of two-color imaging by several groups across many studies (Fig. 1F, see Lu et al., 2016 for review). Even fine-grained details are borne out in these plots. For example, the actin assembly factor Arc15-GFP (a component of the Arp2/3 complex) precedes the actin-binding protein Sac6-GFP at CME sites, and Sac6-GFP is followed closely by Ark1-GFP, a kinase recruited to CME sites by actin binding proteins (Fig. 1F, Cope et al., 1999). Our ability to recapitulate fine-grained differences in timing gave us confidence to draw new conclusions from our dataset.

Maximum intensity and lifetime measurements also recapitulate trends from the literature. While we did not calibrate fluorescence intensity to count molecules, relative fluorescence intensities we measured are in agreement with those reported previously (Fig. S1A, Tables S5-7; Picco et al., 2015; Sun et al., 2019). The median lifetimes we measured are also in general agreement with published results (Fig. S1B, Tables S8-10, Carroll et al., 2012; Kaksonen et al., 2005, 2003; Lu et al., 2016; Peng et al., 2015a; Stimpson et al., 2009). The instances in which our data deviate slightly from the literature are likely due to the exponential decay of TIRFM illumination with distance from the coverslip (Axelrod, 1989). We undercounted Arc15 slightly, most likely due to the actin network extending further from the coverslip than other endocytic components we imaged. Our recorded lifetimes are also systematically slightly lower than those previously reported, likely due to a failure to record the final moments of each CME event when the invagination/vesicle is moving out of the TIRFM field. Despite these minor artifacts, trends within our systematic TIRFM dataset agree with the published literature, empowering our deeper analysis, described below.

One unexpected result from our initial analysis of CME protein dynamics is the amount of variability in our measurements (Fig. 1D-F, Fig. S1A-B). Budding yeast CME has canonically been described as being highly regular, particularly during the late stages (Kaksonen et al., 2006; Stimpson et al., 2009; Sun et al., 2015). However, our dataset made us question this characterization. One measure of variability is the coefficient of variation (CV), the ratio of the standard deviation to the mean (Reed et al., 2002). Our data confirm that the earlier stages of CME (CV: ∼0.5) are more variable then the later stages, but we nevertheless detected considerable variability during the later stages of the process (CV: ∼0.3, See Tables S8-10). Despite repeated characterization of CME as being “highly regular” in the literature, our CV values appear consistent with previously reported data (Kaksonen et al., 2005, 2003; Sun et al., 2006, where CV values can be estimated from plots of mean with standard deviation). We exploited this previously underappreciated variability to gain new systems-level insights into the mechanisms of CME.

### Quantitative evidence for a regulatory transition point in the clathrin-mediated endocytosis pathway

Analysis of paired lifetime and intensity data for individual CME events (as opposed to ensemble averages from many events) revealed at least two distinct behaviors for proteins in the CME pathway. Given the highly interconnected nature of protein interactions at CME sites (Holland and Johnson, 2018; Tonikian et al., 2009), one might predict that longer assembly times at CME sites would lead to greater abundances. To test this prediction, we plotted the maximum intensity from each tracked CME event for each query protein as a function of its lifetime and performed linear regression analyses. The R^2^ value of each fit can be interpreted as the percentage of the variability in maximum intensity that can be explained by the variation in lifetime (Glantz et al., 2001). For early endocytic proteins, exemplified by Ede1, lifetime is poorly correlated with abundance, with linear fits yielding low slopes (0.29) and R^2^ values (0.05) (Fig. 2A, Table S11). Late arriving coat proteins and actin associated proteins, exemplified by Las17, show moderate correlations between lifetime and abundance, with higher slopes (0.97) and better fits (R^2^ = 0.21) (Fig. 2B, Table S11). Finally, proteins involved in scission and disassembly of the endocytic machinery, exemplified by Rvs167, once again have abundances that are poorly predicted by lifetimes (slope = 0.37, R^2^ = 0.08, Fig. 2C, Table S11). To summarize the results from our analysis of each CME protein imaged, we generated bar graphs with the height of the bars representing the slope of the fit and the color of the bars representing the R^2^ value (Fig. 2D, see Fig. S2 for individual plots, Table S11 for fit statistics). From these plots, two behaviors for CME proteins emerge. Early arriving proteins and very late arriving proteins have abundances that are largely independent of their lifetimes, while components of the endocytic coat and WASP/Myosin module show a stronger correlation between these parameters. Such temporally resolved differences in behavior hint that a shift in CME protein behavior may occur sometime after endocytic site initiation but prior to actin assembly.

**Figure 2:**
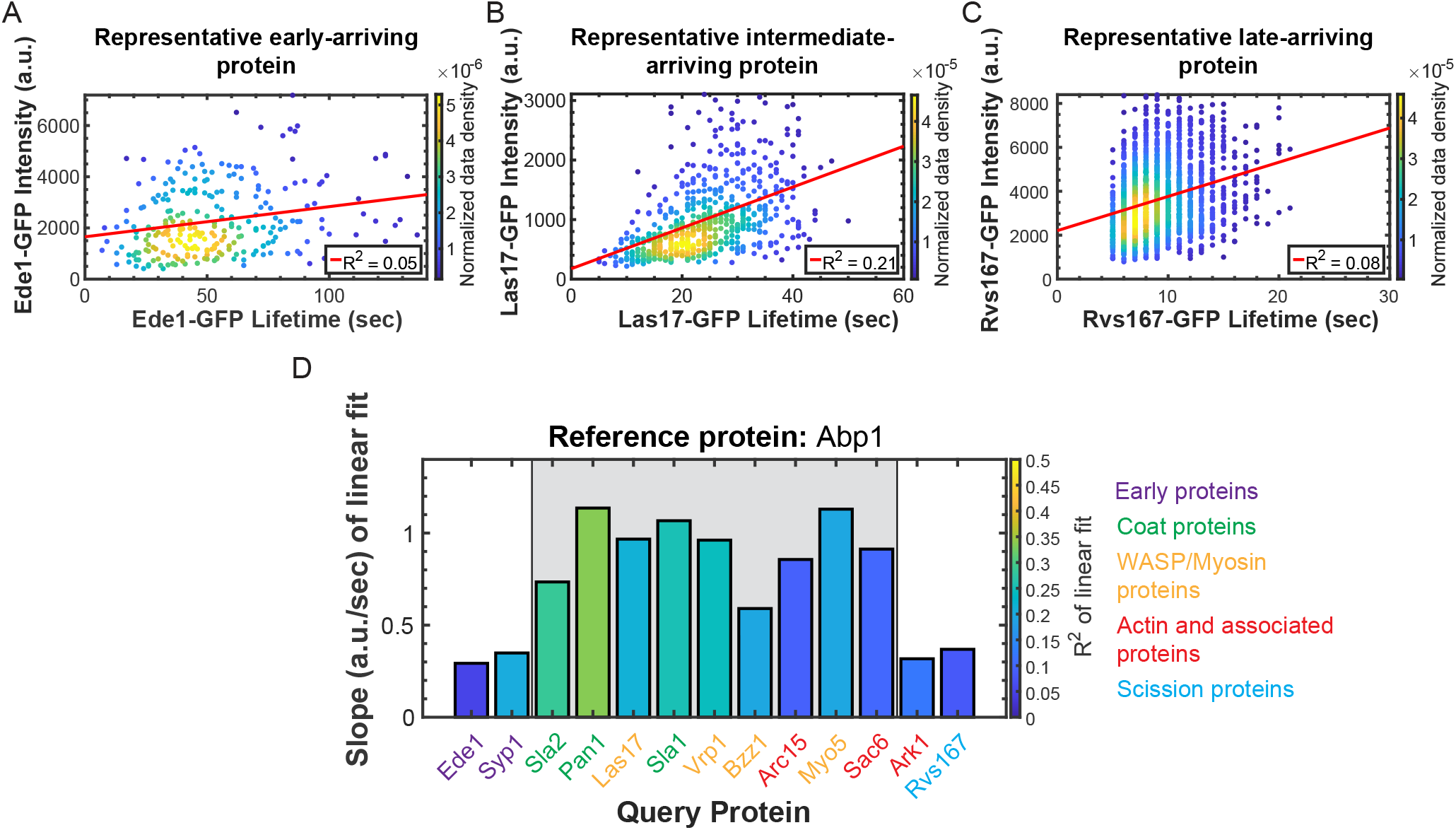
Lifetime and maximum intensity are positively correlated for individual later arriving CME proteins, but are less correlated for earlier arriving proteins. (A – C) Scatterplots of maximum intensity vs. lifetime for (A) Ede1-GFP, (B) Las17-GFP, and (C) Rvs167-GFP. Color indicates the normalized data density in the neighborhood around each point. Red lines are linear fits to the data with the indicated R^2^ values. (D) Summary of linear fits to plots of maximum intensity vs. lifetime for the indicated GFP-tagged query proteins. The bar height and color indicate the normalized slope and the R^2^ of the fit, respectively. The grayed area highlights proteins displaying similar behavior. Colored text indicates which module each protein imaged belongs to according to the color scheme at right. Association of query protein signal with signal of the reference protein Abp1-RFP was used to eliminate spurious events. See Fig. S2 for all related scatter plots.

We next investigated interrelationships between the behaviors of different CME proteins. Many previous studies have presented two-color imaging of CME proteins (Kaksonen et al., 2003, 2005). However, previous analyses have typically relied on separating and averaging measurements of the two protein species imaged, so correlations between the behaviors of the two proteins at individual CME events imaged were lost (e.g., Kaksonen et al., 2005).

Since some CME proteins have lifetimes that are correlated with their own maximum intensities, and because CME proteins are extensively interconnected through protein-protein interactions, we wondered whether the lifetimes of any of our query proteins would be predictive of the abundances of our reference proteins. When we plotted the maximum intensity of our reference proteins at each CME event as a function of the lifetime of the corresponding query protein, we once again detected a quantitative transition point within the CME pathway. Early proteins such as Ede1 have lifetimes that are poorly correlated with the maximum abundance of the CME actin marker Abp1 (slope = −0.06, R^2^ = 0.00, Fig. 3A, Table S12). In contrast, lifetimes of late arriving coat proteins, WASP/Myosin module proteins, and actin binding proteins, exemplified by Las17, are moderately predictive of maximum Abp1 levels (slope = 0.96, R^2^ = 0.13, Fig. 3B, Table S12). Finally, lifetimes of proteins involved in scission and disassembly, exemplified by Rvs167, are once again less correlated with maximum Abp1 intensity (slope = 0.28, R^2^ = 0.04, Fig. 3C, Table S12). We again summarized the data from our linear fits by making bar graphs as in Fig. 2D (Fig. 3D, see Fig. S3 for individual plots, Table S12 for fit statistics). While the lifetimes of the earliest and latest arriving CME proteins are poorly correlated with the abundance of the actin marker Abp1, all proteins of the late coat and WASP/Myosin CME modules have lifetimes that are comparatively more positively correlated with Abp1 abundance (Fig. 3D). These data show that the extent of actin assembly is independent of the behavior of early arriving proteins, but more dependent on the behavior of later arriving proteins, consistent with what we know about the interaction network for CME proteins. They imply the existence of a transition point in the pathway separating early proteins from later ones.

**Figure 3:**
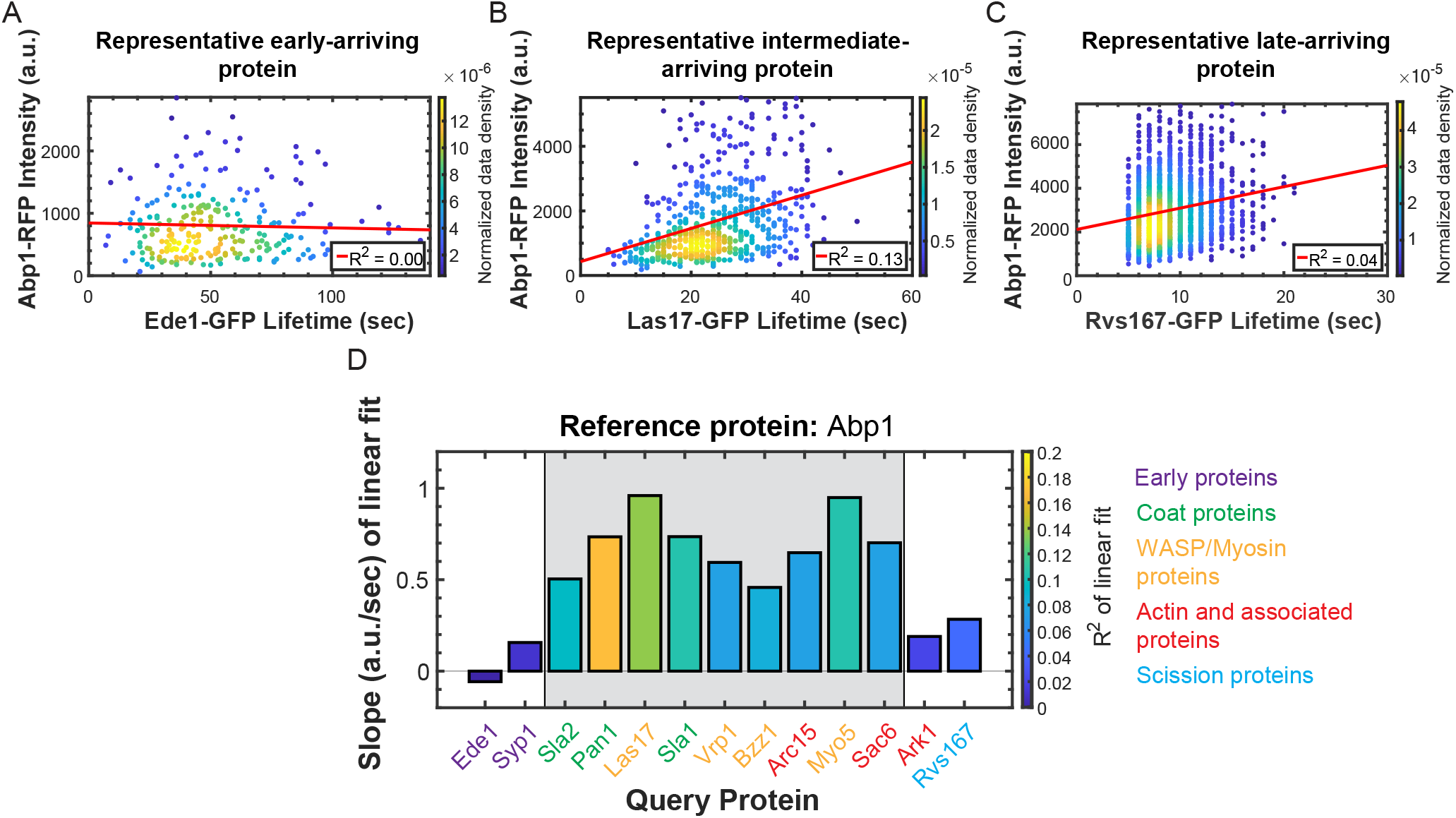
Correlations between reference protein abundance and query protein lifetime provide further evidence for a regulatory transition point. (A – C) Scatterplots of the maximum intensities of Abp1-RFP vs. the lifetimes of (A) Ede1-GFP, (B) Las17-GFP, and (C) Rvs167-GFP as in Fig. 2A-C. (D) Summary of linear fits to plots of maximum Abp1-RFP intensity vs. lifetime for the indicated GFP-tagged query proteins as in Fig. 2D. See Fig. S3 for all related scatter plots.

To determine how behaviors of individual CME proteins are correlated with progression through the pathway as a whole, we tested whether lifetimes of any of our query proteins are predictive of the lifetimes of our reference proteins. Early proteins, once again exemplified by Ede1, have lifetimes that are well correlated with other early protein lifetimes (slopes ranging from 0.45 – 0.71, R^2^ values ranging from 0.21 – 0.55), but poorly correlated with the lifetimes of the late arriving proteins Sla1 and Abp1 (slopes of 0.17 and 0.08 respectively, R^2^ values of 0.06 and 0.03 respectively, Fig. 4A, Tables S13-15). Late arriving coat proteins and proteins of the WASP/Myosin module have lifetimes that are moderately positively correlated to the lifetime of actin assembly as reported by Abp1 (for Las17, slope = 0.4 and R^2^ = 0.24, Fig. 4B, Table S15). Actin module proteins, such as Sac6, have lifetimes that are well correlated with the lifetime of the actin marker Abp1 (slope = 0.83, R^2^ = 0.71, Fig. S5, Table S15). Finally, scission module proteins have lifetimes that are moderately well correlated with Abp1 lifetime (for Rvs167, slope = 0.21, R^2^ = 0.15, Fig. 4C, Table S15). All queried early arriving proteins have lifetimes that are well correlated with the lifetime of the early arriving reference protein Ede1 (Fig. 4D, Fig. S4, Table S13), suggesting that a signal or a stochastic event, the nature of which is currently unknown, may be required to trigger concerted maturation of the early site. However, long lifetimes of early arriving CME proteins are not predictive of the lifetime of any of the later-arriving reference proteins that we tested; our queried early CME proteins have lifetimes that are poorly correlated with both Sla1 and Abp1 lifetimes (Fig. 4E-F, Fig. S4-5, Tables S14-15). Thus, while behavior of any one early arriving CME protein is predictive of the behavior of the others, the early proteins are molecularly insulated from the proteins in the late part of the CME pathway.

**Figure 4:**
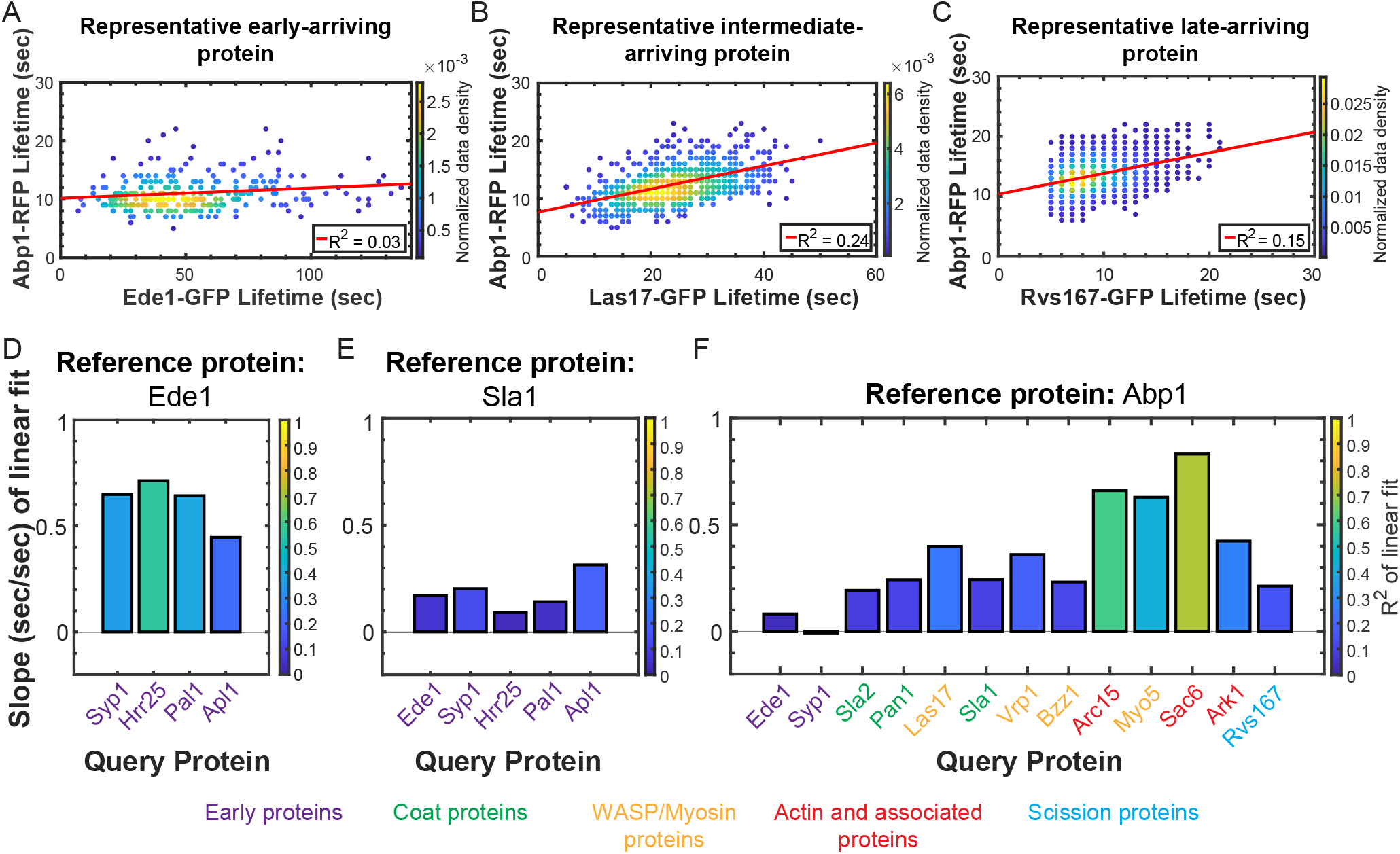
CME protein lifetimes are correlated amongst earlier arriving proteins, but early protein lifetimes are not correlated with late protein lifetimes. (A – C) Scatterplots of the lifetimes of Abp1-RFP vs. the lifetimes of (A) Ede1-GFP, (B) Las17-GFP, and (C) Rvs167-GFP as in Fig. 2A-C. (D – F) Summaries of linear fits to plots of (C) Ede1-RFP, (E) Sla1-mCherry, and (F) Abp1-RFP lifetime vs. lifetime for the indicated GFP-tagged query proteins as in Fig. 2D. See Figs. S1 - 3 for all related scatter plots.

Together, these data indicate that the CME pathway in budding yeast can be thought of as occurring in two, quantitatively recognizable phases. Early arriving proteins initiate CME sites and persist for a variable period of time, but the length of this variable phase is not predictive of events that occur downstream in the pathway. Around the time of Sla2 recruitment, the CME site transitions into a more regular phase. The behavior of any one protein predicts the behavior of other proteins in this phase, although there is still variability in the behavior of proteins. Sla2 has been called an intermediate coat protein (Carroll et al., 2012), and it is indeed “intermediate” by nearly every measure in our data set. Its lifetime is less variable than those of the early proteins, but more variable than the lifetimes of the late phase proteins (Early protein CV ∼0.5, Late protein CV ∼ 0.3, Sla2 CV ∼0.4. Tables S8-10). The lifetime of Sla2 is also intermediately correlated with its own abundance, the abundance of Abp1, and the lifetime of Abp1 (Fig. 2D, Fig. 3D, and Fig. 4F). Sla2 may therefore be involved in transitioning from the early phase of CME into the late phase. Having substantiated the existence of this transition point based on an unbiased quantitative imaging approach, we decided to investigate molecular determinants that dictate the rate of maturation from the early phase into the late phase.

### Maturation through the CME transition point is faster in cellular regions with concentrated endocytic cargo

Cargo has previously been suggested to influence the maturation rate of CME sites (Carroll et al., 2012; Layton et al., 2011). We set out to determine whether cargo influences maturation through tuning the rate of transition from the early phase of CME to the late phase.

As an initial strategy for varying the presence of cargo at CME sites, we took advantage of polarized secretion during the budding yeast cell cycle. When mitotically replicating yeast are in the small to medium-budded cell cycle stages, exocytic events are localized primarily to the growing bud, whereas exocytic events are redirected toward both sides of the bud neck at cytokinesis (Lew and Reed, 1995). Endocytic cargos such as vesicle-associated SNARE proteins must be retrieved from the plasma membrane by CME following exocytosis (Lewis et al., 2000). Because septin filaments at the bud neck prevent lateral diffusion between the mother and the bud (Takizawa et al., 2000), the polarized pattern of secretion in small-budded cells concentrates CME cargo in the bud, creating cargo-poor mothers, whereas large-budded cells have more cargo delivered to the mother cell plasma membrane.

To determine whether the presence of endocytic cargo influences the rate of transition from the early phase of CME to the late phase, we imaged a representative protein from each respective endocytic phase through the cell cycle. We chose Ede1-GFP as a marker of the early phase of CME and Sla1-mCherry as a marker of the late phase. During the small- and medium-budded stages of the cell cycle, when cargo is plentiful in the bud but sparse in the mother, we observed considerable polarization of Sla1-mCherry-marked late CME sites, while Ede1-GFP-marked early CME sites were notably less polarized (Fig. 5A, Video 2). The disparity in polarization diminishes as cells near cytokinesis, when cargoes are more evenly distributed between the mother and bud (Fig. 5A, Video 2). As cells begin the next cell cycle, Sla1-marked late phase CME sites again become more polarized (Fig. 5A, Video 2). Based on the observation that early CME sites were not as remarkably polarized into buds as late sites within the same cell, we concluded that the presence of cargo does not appreciably affect the rate of CME site initiation as has been previously proposed (Godlee and Kaksonen, 2013; Goode et al., 2015), but that it increases the proportion of initiated CME sites that mature.

**Figure 5:**
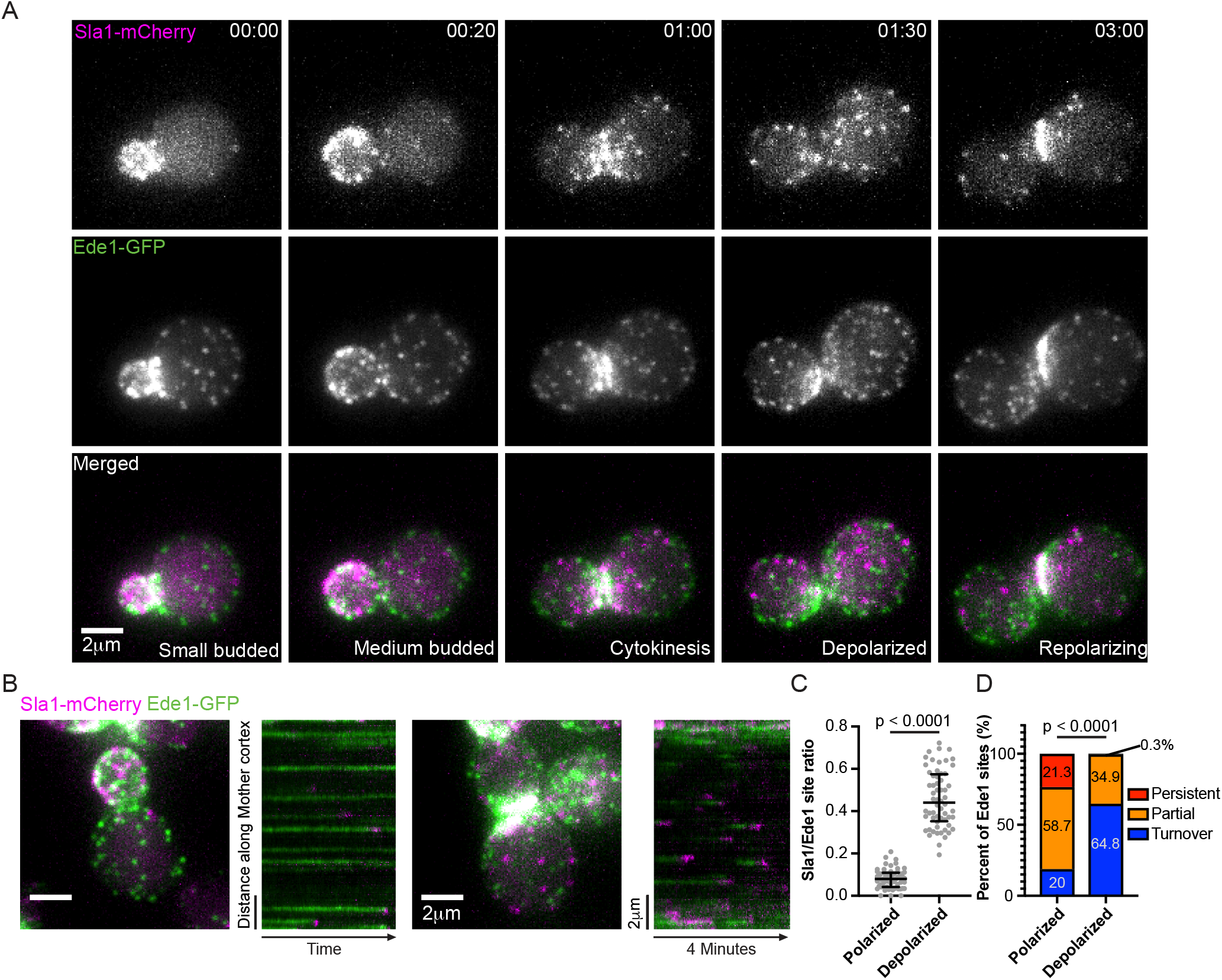
Proximity to sites of exocytosis and cell polarity signaling influences the rate of endocytic site maturation. (A) Montage from a maximum intensity-projected epifluorescence video of a cell endogenously expressing Sla1-mCherry (magenta) and Ede1-GFP (green). Times are hours:minutes. See Video 2. (B) Maximum intensity projections of z-stacks (left) of polarized and depolarized cells paired with circumferential kymographs around the mother cortex from medial focal plane videos of the same cells (right) endogenously expressing Sla1-mCherry (magenta) and Ede1-GFP (green). See Video 3. (C) Quantification of the ratio of the number of Sla1-mCherry sites to Ede1-GFP sites in mother cells from maximum intensity projections of z-stacks of 60 polarized and 60 depolarized cells. A two-tailed p value from a Mann-Whitney U test with the null hypothesis that the two ratios are identical (U = 1) is displayed. The median and interquartile ranges are denoted with error bars. (D) Percentage of 596 Ede1 patches from 59 polarized mother cells and 1034 Ede1 patches from 56 depolarized mother cells that persist throughout the duration of a 4-minute video (persistent), that are present at the start or end of the video (partial), or that assemble and disassemble within the interval of the video (turnover). Numbers indicate the percentage of patches observed in each category. A two-tailed p value from a Chi-Square test with the null hypothesis that the proportion distributions are identical (chi-square = 1349, 2 degrees of freedom) is displayed.

Because all CME sites appear to follow the same molecular pathway in budding yeast (Kaksonen et al., 2003), the abundance of early CME sites relative to late CME sites in small-budded mother cells could be explained by the presence of either abortive or stalled CME events in these mother cells. The “extra” Ede1-GFP-marked early sites that we observed could either abort, leading to the dearth of Sla1-marked late CME sites in the mother cells, or they could mature at a slower rate, also resulting in fewer late phase sites. To distinguish between these possibilities, we used a two-step imaging regime to first identify highly polarized and depolarized cells, and to then track the fate of early CME sites in mother cells of each cell type. First, we collected 5 µm z-stacks (11, 500 nm slices) to identify highly polarized cells, defined as small-budded cells with 3 or fewer Sla1-mCherry-marked late CME sites in the mother, and depolarized cells, defined as medium- or large-budded cells with greater than 3 Sla1-mCherry-marked late CME sites in the mother (Fig. 5B, left photos). Next, we collected time series in the medial focal plane of the same cells, capturing images every 2 seconds for 4 minutes, and generated circumferential kymographs along the mother cortex (Fig. 5B, right photos, Video 3). Our imaging protocol allowed us to confidently identify highly polarized and depolarized cells and to visualize CME at high temporal resolution in the cells identified.

When we analyzed the fates of individual CME sites in mother cells, we determined that the rate of maturation through the transition point is faster when cells are depolarized, i.e., when exocytosis shows no preference for the mother cell or the bud. Quantification of the ratio of the number of late CME sites to early sites in mothers of highly polarized cells and mothers of depolarized cells confirms the observation that there are significantly more Ede1-GFP-marked early CME sites per Sla1-mCherry-marked late site in small-budded mother cells (median ratio of 0.08 vs. 0.44, Fig. 5B left photos, Fig. 5C). Tracking the fate of every early CME site in the medial focal plane in circumferential kymographs failed to reveal any abortive CME events, but it revealed many persistent early sites (Fig. 5B right photos, Fig. 5D, Video 3). While we could not reliably extract Ede1 lifetimes due to a large fraction of Ede1 puncta being present at the start and end of our movies, we observed significantly more long-lived Ede1-marked early CME sites in kymographs from mothers in highly polarized cells than from mothers in depolarized cells (21.3% persistent sites compared to 0.3%, 58.7% “partial” sites compared to 34.9%, Fig. 5D, see figure legend). This observation suggests that the maturation rate from the early phase of CME to the late phase is slower when CME sites are far from cargo and the cell polarity machinery, as they are in small-budded mothers, when the majority of cargo is delivered to the bud. When we conducted the same kinds of experiments with two late-phase CME markers, Sla1-GFP and Abp1-RFP, neither the ratio of numbers of puncta nor the lifetimes of these CME markers were sensitive to polarization state (median ratios 0.56 compared to 0.63, Fig. S6A-C). While Abp1 lifetimes differ slightly (by 0.5 s, p = 0.0247), the small magnitude of this difference makes it unlikely to be biologically relevant. Based on these observations, we concluded that proximity to cargo and/or polarity machinery accelerates the rate of maturation from the early phase of CME to the late phase, but does not affect the kinetics of late events in CME or the CME site initiation rate.

### The delivery of cargo, rather than cell cycle stage, dictates maturation rate through the regulatory transition point

To determine whether the differences we observed in CME site maturation rate were due to polarization state or cell cycle stage, we used osmotic shock to reposition CME cargos and cell polarity proteins independent of cell cycle stage. A previous study from our lab used a temperature-sensitive secretion mutant to halt accumulation of endocytic cargo on the plasma membrane and reported increased numbers of long-lived Ede1-GFP-marked CME sites (Carroll et al., 2012). We conducted the converse experiment by redistributing endocytic cargo from the plasma membrane of buds in highly polarized cells to the plasma membrane of mother cells, testing whether long-lived early phase CME sites could be induced to mature through loss of cell polarity and redirection of secretion. Diluting *S. cerevisiae* cells out of media into water has been reported to trigger depolarization and a wave of CME that internalizes diverse surface proteins, making this treatment a convenient way of manipulating CME cargo (Pringle et al., 1989; Lang et al., 2014). We found that osmotically shocking highly polarized cells by rapidly diluting them with water caused dispersal of the polarity marker Bem1-GFP (Fig. 6A-B). This treatment also results in redistribution of GFP-Sec4-marked secretory vesicles from a bud-localized distribution to a more uniform one, suggesting that exocytic traffic (and therefore endocytic cargo) is repositioned by this manipulation (Fig. S6D). We therefore used osmotic shock to disperse cargo independent of the cell cycle.

**Figure 6:**
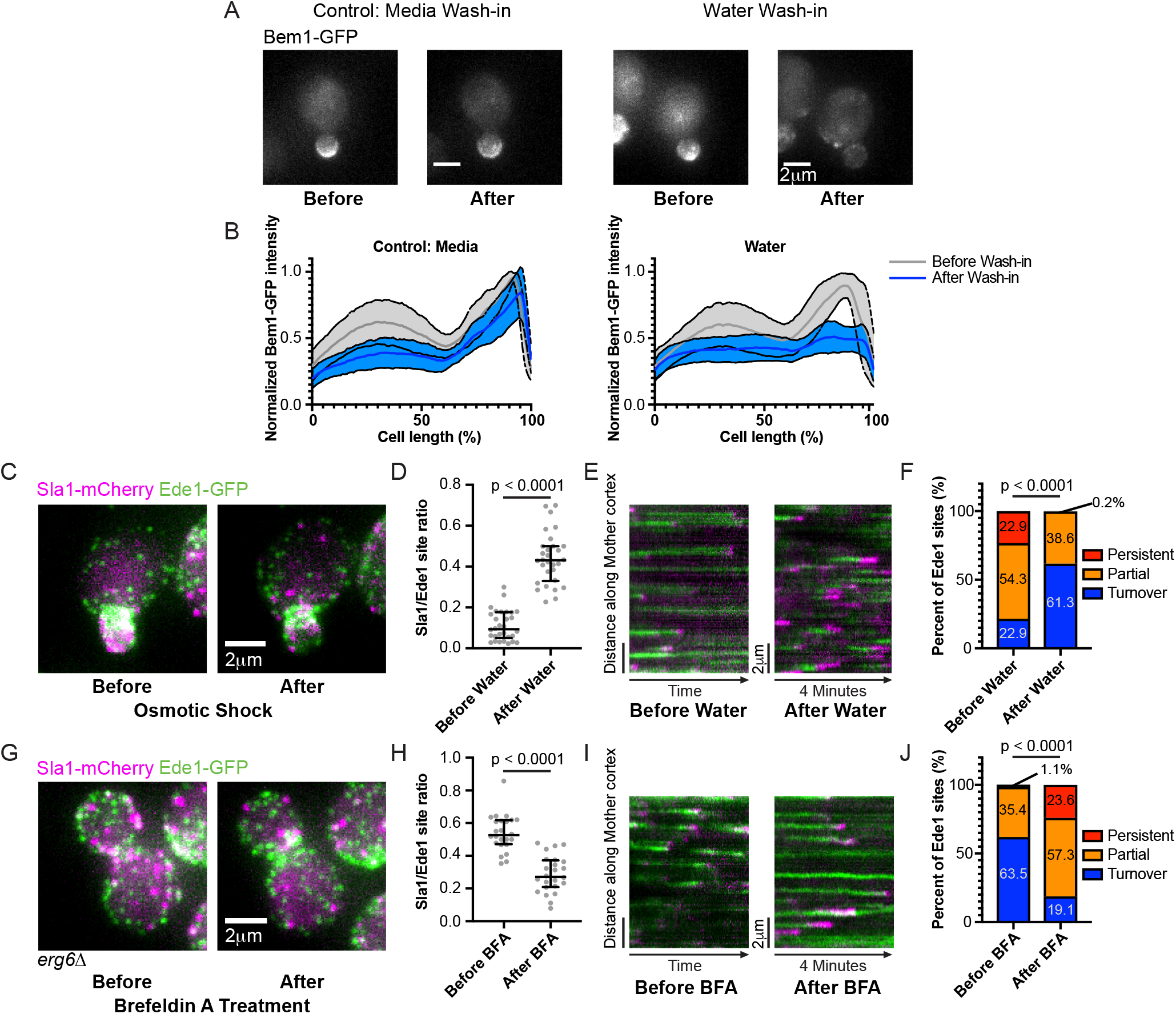
Endocytic site maturation rate is controlled by polarized cargo deposition. (A) Maximum intensity projections of cells endogenously expressing polarity marker Bem1-GFP before and 5 minutes after 17-fold dilution into isotonic imaging media (control, left) or water (right). (B) Average ± standard deviation for Bem1-GFP intensity profiles from 20 (media, left) and 30 (water, right) cells before and after dilution into the indicated media. Individual intensity profiles generated from 25 pixel-wide lines were normalized to the maximum value before dilution and to cell length. (C) Maximum intensity projections of a cell endogenously expressing Sla1-mCherry (magenta) and Ede1-GFP (green) before and 5 minutes after 17-fold dilution into water. (D) Quantification of the ratio of the number of Sla1-mCherry sites to Ede1-GFP sites from maximum intensity projections of z-stacks of 30 small-budded mother cells before and after osmotic shock. A two-tailed p value from a Mann-Whitney U test as in Fig. 5C (U = 7) is displayed. The median and interquartile ranges are denoted with error bars. (E) Circumferential kymographs around the mother cortex from medial focal plane videos of cells endogenously expressing Sla1-mCherry (magenta) and Ede1-GFP (green) before and 5 minutes after 17-fold dilution into water. (F) Percentage of 280 Ede1 patches from 30 small-budded mother cells before and 594 Ede1 patches from 30 small-budded mother cells after osmotic shock that persist, are partially captured, or turnover during a 4-minute video as in Fig. 5D. A two-tailed p value from a Chi-Square test as in Fig. 5D (chi-square = 543.1, 2 degrees of freedom) is displayed. (G) Maximum intensity projections of an *erg6Δ* cell endogenously expressing Sla1-mCherry (magenta) and Ede1-GFP (green) before and 10 minutes after treatment with Brefeldin A. (H) Quantification of the ratio of the number of Sla1-mCherry sites to Ede1-GFP sites from maximum intensity projections of z-stacks of 24 large-budded, *erg6Δ* mother cells before and after BFA treatment. A two-tailed p value from a Mann-Whitney U test as in Fig. 5C (U = 25.5) is displayed. The median and interquartile ranges are denoted with error bars. (I) Circumferential kymographs around the mother cortex from medial focal plane videos of *erg6Δ* cells endogenously expressing Sla1-mCherry (magenta) and Ede1-GFP (green) before and 10 minutes after treatment with BFA. (J) Percentage of 277 Ede1 patches from 18 large-budded, *erg6Δ* mother cells before and 266 Ede1 patches from 22 large-budded, *erg6Δ* mother cells after BFA treatment that persist, are partially captured, or turnover during a 4-minute video as in Fig. 5D. A two-tailed p value from a Chi-Square test as in Fig. 5D (chi-square = 1122, 2 degrees of freedom) is displayed.

Forced depolarization through osmotic shock accelerates maturation through the transition point in small-budded mother cells. When we examined the CME markers Ede1-GFP and Sla1-mCherry in small-budded mothers before and after osmotic shock, we found that the treatment caused Sla1-mCherry sites to depolarize. Ede1-GFP sites were not affected (Fig. 6C). Quantification of the ratio of the number of late phase CME sites to early phase CME sites in these mother cells reveals a change in ratio reminiscent of the one that occurs naturally during the cell cycle (median ratio changes from 0.09 to 0.43, Fig. 6D, see also Fig. 5C). To determine whether this transition was caused by a change in the maturation rate of CME sites, we examined Ede1-GFP tracks in circumferential kymographs from mother cells before and after osmotic shock (Fig. 6E). We observed a significant decrease in the proportion of long-lived early CME sites upon osmotic shock, suggesting that early phase CME sites mature to the late, regular phase more quickly (from 22.9% persistent sites to 0.2%, 54.3% partial sites compared to 38.6%, Fig. 6F). Interestingly, the shortening of Ede1-GFP lifetimes by osmotic shock does not extend to late phase CME proteins. While the ratio of the number of Abp1-RFP to Sla1-GFP puncta was unchanged, both Sla1-GFP and Abp1-RFP lifetimes were significantly increased upon osmotic shock (median lifetimes of 22 and 10 s, respectively, before osmotic shock and 34 and 13 s, respectively, after. Fig. S6E-H). Lengthening of Sla1-GFP and Abp1-RFP lifetimes is likely due to a mechanical burden on membrane invagination/internalization caused by increased turgor pressure in hypotonic solution (Hassinger et al., 2017; Aghamohammadzadeh and Ayscough, 2009; Boulant et al., 2011). Our osmotic shock experiments suggest that moving endocytic cargo from places where it is concentrated to places where it is scarce can activate stalled endocytic sites, even as the internalization phase is slowed.

In contrast to forced depolarization, elimination of endocytic cargo causes endocytic sites to stall. We acutely blocked accumulation of endocytic cargo on the plasma membrane through treating cells with the drug Brefeldin A (BFA), a secretion pathway inhibitor (Graham et al., 1993). BFA inhibits secretion in cells lacking the sterol synthesis gene *ERG6*, but it does not affect secretion in wild-type cells (Graham et al., 1993). Large-budded *erg6Δ* cells resemble large-budded wild-type cells, with both early and late CME sites visible in the mother (Fig. 6G, Fig. S6I). Treating *erg6Δ* cells with BFA significantly reduced the ratio of late endocytic sites to early endocytic sites in large-budded mothers (Fig. 6G-H). Kymographs along the mother cortex and analysis of early endocytic site turnover indicate that slowed maturation rate and increased frequency of persistent early endocytic sites accounts for this change in ratio of late endocytic sites to early ones (Fig. 6I-J). Interestingly, treatment of wild-type cells with BFA caused a slight increase in the ratio of late endocytic sites to early ones, accounted for by a slight drop in the proportion of endocytic sites that turnover, although this effect was very small (Fig. S6I-L). Although each approach we used (classification of cells by cell cycle state, osmotic shock, and BFA treatment) reflects or results in changes to multiple cellular processes, the common denominator between them is an effect on cargo availability. Together, these data demonstrate that availability of cargo at CME sites licenses maturation from the early phase to the late phase of the pathway. The transition point we identified can therefore be considered a cargo checkpoint.

### The regulatory transition point occurs between recruitment of the intermediate coat proteins and late coat proteins

Since a representative CME protein from before the regulatory transition point (Ede1) was not significantly polarized in small-budded cells but a late phase CME protein (Sla1) was (Fig. 5), we used this difference in CME protein distribution as an assay to pinpoint the time of the regulatory transition point in the CME pathway. We recorded Z-stacks of proteins from the early, early coat, intermediate coat, and late coat modules using either epifluorescence microscopy or spinning disc microscopy (depending on how well we could distinguish CME puncta above background noise). The early proteins Syp1 and Hrr25, the early coat proteins Apl1 and Pal1, and the intermediate coat proteins Sla2, Ent1, and Ent2 all display distributions similar to that of Ede1: in highly polarized mother cells with few Sla1-mCherry-marked late CME sites, we nevertheless observed many puncta of these proteins, placing them upstream of the transition point (median ratios of 0.14 vs. 0.59 (Syp1), 0.09 vs. 0.49 (Sla2), 0.16 vs. 0.79 (Ent2), Fig. 7A-B, Fig. S7A). The late coat marker Pan1, on the other hand, polarizes with Sla1-mCherry-marked late CME sites (median ratio of 0.62 vs. 0.64, Fig. 7A-B). These data indicate that the transition point in the CME pathway occurs after recruitment of the intermediate coat module but before recruitment of the late coat proteins Pan1 and Sla1.

**Figure 7:**
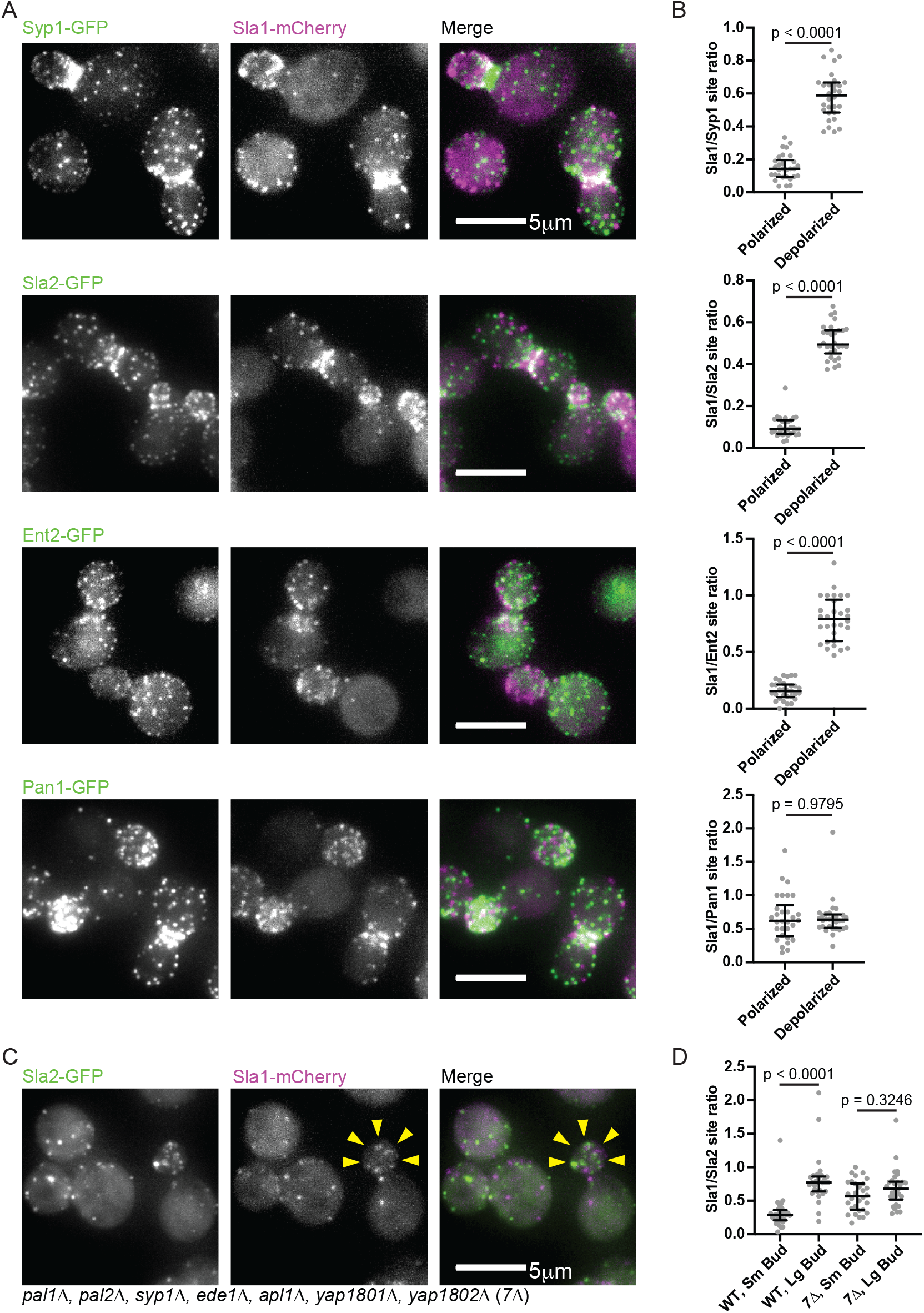
Defining molecular components of the early and late steps of the endocytic pathway. (A) Maximum intensity projections of epifluorescence z-stacks of clusters of cells endogenously expressing Sla1-mCherry (magenta) and Syp1-GFP (green, first row), Sla2-GFP (green, second row), Ent2-GFP (green, third row), and Pan1-GFP (green, last row). Individual channels are shown in gray scale at left. (B) Quantification of the ratio of the number of Sla1-mCherry sites to the number of Syp1-GFP (top row), Sla2-GFP (second row), Ent2-GFP (third row), and Pan1-GFP (last row) sites from maximum intensity projections of z-stacks of small-budded (polarized) and large-budded (depolarized) mother cells (n = 30 cells per category). Two-tailed p values from Mann-Whitney U tests as in Fig. 5C (U_Syp1_ = 0, U_Sla2_ = 0, U_Ent2_ = 0, U_Pan1_ = 448) are displayed. The median and interquartile ranges are denoted with error bars. (C) Maximum intensity projections of epifluorescence z-stacks of clusters of *7Δ* cells endogenously expressing Sla1-mCherry (magenta) and Sla2-GFP (green). Individual channels are shown in gray scale at left. Yellow arrowheads indicate highlight low concentration of Sla1-mCherry sites in a small bud, where late endocytic sites would normally be plentiful. (D) Quantification of the ratio of the number of Sla1-mCherry sites to the number of Sla2-GFP sites in small budded and large budded wild-type and *7Δ* cells (n = 30 cells per category). Numbers are p values from Kruskal-Wallis tests followed by Dunn’s multiple comparisons test. The median and interquartile ranges are denoted with error bars.

Since we concluded that polarized cargo distribution drives polarization of late endocytic sites during the cell cycle through controlling maturation rate, we predicted that mutant cells that could not sense cargo at endocytic sites would also fail to undergo cycles of late endocytic site polarization and depolarization. A mutant with genes encoding seven early arriving endocytic proteins deleted (“*7Δ*”) has previously been reported to complete endocytosis without internalizing cargo (Brach et al., 2014). When we examined these mutants, we found that they were no longer polarized during the small-budded stage of the cell cycle (Fig. 7C). We compared the ratio of late CME sites (marked by Sla1-mCherry) to early sites (marked by Sla2-GFP) in small- and large-budded *7Δ* mothers and found that the ratio was no longer significantly different in the *7Δ*mutants (Fig. 7 C-D). In contrast, small- and large-budded mothers were insignificantly different with respect to the ratio of the number of puncta of two late endocytic markers to one another (Fig. S7B-C). Thus the cargo-sensitive maturation rate of individual endocytic sites accounts for polarization of late endocytic sites during specific phases of the cell cycle.

## Discussion

Despite identification of dozens of proteins involved in CME through decades of research, how progression through this pathway is regulated is poorly understood. In this study, we conducted systematic imaging experiments of CME in budding yeast to reveal new regulatory principles. Previous studies used two-color imaging to describe a protein recruitment cascade at CME sites; however, these studies typically analyzed the recruitment dynamics of each protein independently, therefore losing information about how the behavior of one protein is correlated with the behavior of another at individual endocytic sites. To uncover inter-relationships in the behavior of individual components of the CME machinery, we performed quantitative analysis of the pair-wise recruitment and abundance behavior of CME proteins at >17,000 endocytic sites.

Our data set revealed more variation in the behavior of individual proteins in the budding yeast CME pathway than had previously been appreciated. CME has been described as a highly regular molecular pathway. The ordered arrival of CME proteins at endocytic sites is roughly conserved from yeasts to humans and is nearly invariant (Kaksonen et al., 2005, 2003; Taylor et al., 2011; Doyon et al., 2011; Sun et al., 2019). While the initial steps of CME were known to occur with variable timing, the timing of the late steps of the pathway had been considered to be regular (Kaksonen et al., 2006). Previous studies even used lifetimes of late phase CME proteins as readouts for endocytic efficiency (Kaksonen et al., 2005). While our data confirm that the early stages of CME occur with more variable timing than the late stages, we were surprised to also observe variability in what has previously been considered the “regular phase” of the process. We took advantage of this previously underappreciated variability in timing to gain new insights into the CME pathway by looking for covariation in the pair-wise behavior of endocytic proteins.

Our analysis suggests that a full complement of early arriving CME proteins is present at each endocytic site at initiation. Each early arriving endocytic protein has been reported to have a variable lifetime (Carroll et al., 2012, 2009; Peng et al., 2015; Stimpson et al., 2009; Newpher et al., 2005). However, the implication of this variability was unknown. One possible interpretation was that long-lived early CME sites are incompletely assembled, potentially lacking some component of the early or early coat module. Another possibility was that the early proteins assemble fully but that maturation to the next phase of the pathway is stalled due to an inhibitory checkpoint signal. Our pair-wise analysis of the lifetimes of each early arriving protein and Ede1 shows that they are well correlated. This correlation is consistent with the possibility that CME proteins assemble at the nascent sites in a coordinated manner, but that advancement to the next phase awaits release of a checkpoint.

Analyses of correlations between the behaviors of different pairs of proteins at individual CME sites also reveal two quantitatively separable stages, substantiating the existence of a regulatory transition point in the CME pathway (Carroll et al., 2012, Fig. 8A). This conclusion is based on the loss of correlation of lifetimes and intensities for proteins spanning a specific step in the pathway. While lifetimes of early arriving proteins are predictive of one another (for example, a plot of Ede1 lifetime vs. Syp1 lifetime yields a linear fit with a slope of 0.65 and an R^2^ of 0.35), in no case does the behavior of an early arriving protein strongly correlate with the behavior of a late arriving protein (for example, a plot of Ede1 lifetime vs. Abp1 lifetime yields a linear fit with a slope of 0.08 and an R^2^ of 0.03). Conversely, lifetimes of late arriving proteins are fairly good predictors of the maximum intensity of the endocytic actin reporter Abp1 (Slopes ranging from 0.46 to 0.96). The striking correlations within groups of proteins recruited early or late, but not between these groups, is consistent with the notion that robust protein interaction networks are at play during CME: an early complex of proteins establishes endocytic sites and a late complex carries out membrane invagination and vesicle scission (Fig. 8A). The poor correlation between the lifetimes and intensities of these two networks likely indicates that there are fewer molecular links between the two networks than within each network. Since proteins in the early and late phases have behaviors that are predictive of behaviors of other proteins in the same phase, but not the other phase, CME initiation and internalization can be thought of as two separable processes.

**Figure 8:**
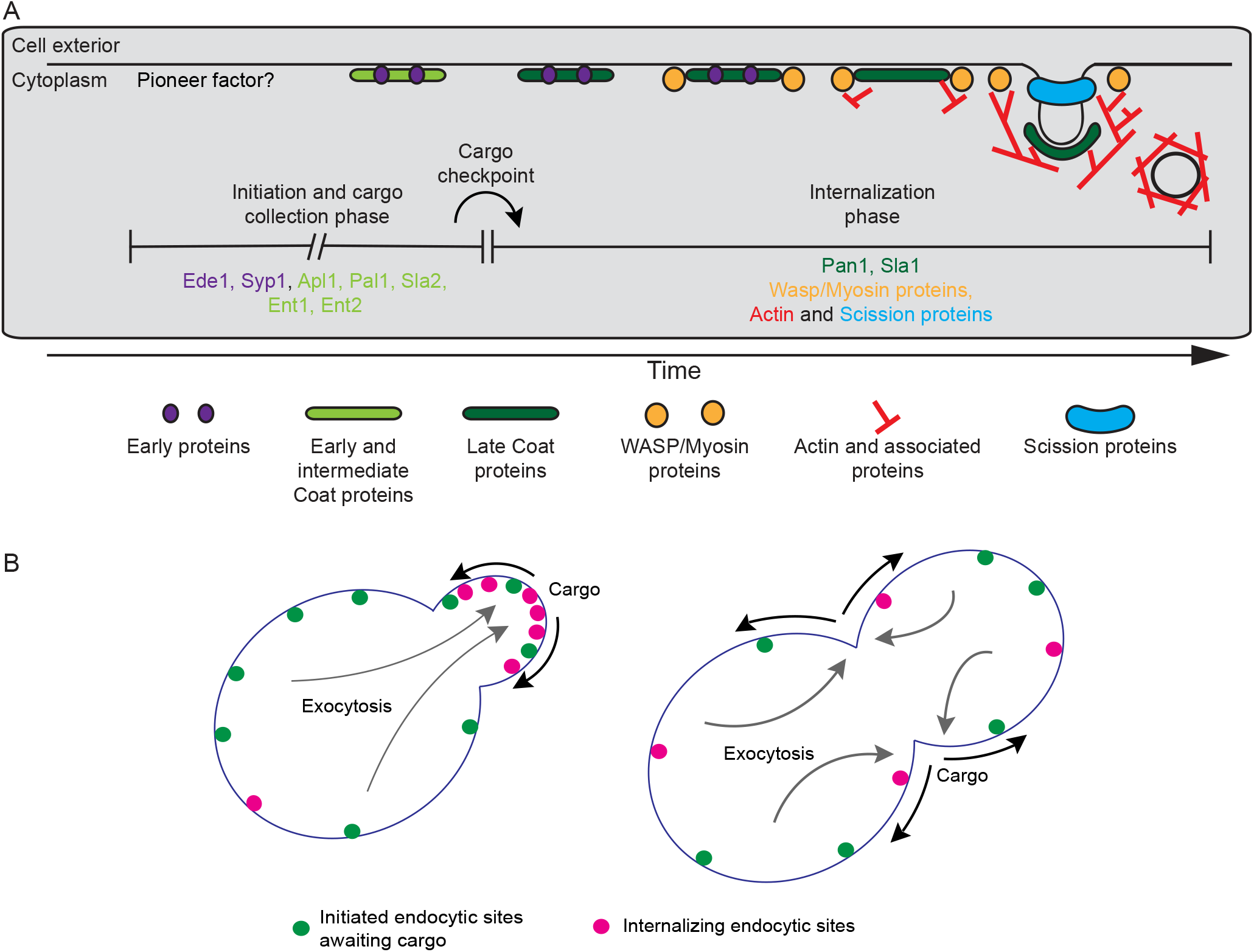
Cargo regulates the transition between two quantitatively distinguishable phases of endocytosis. (A) Revised timeline for the clathrin-mediated endocytosis pathway indicating the proposed temporal location of the cargo checkpoint. Proteins upstream and downstream of the checkpoint are listed under each phase. (B) Schematic cartoon illustrating the proposed mechanism leading to polarized endocytosis in budding yeast. In highly polarized cells, endocytic cargos (black arrows) are delivered primarily to the growing bud by exocytosis (gray arrows), locally accelerating the transition from the initiation and cargo-collection phase of CME (green spots) into the internalization phase (magenta spots). As exocytosis becomes depolarized later in the cell cycle, cargos are more plentiful across the plasma membrane, leading to global acceleration of maturation through the cargo checkpoint.

Through observation of CME site composition and dynamics while directionality of polarized yeast growth changed naturally in a programmed manner through the cell cycle, and through genetic and chemical perturbation of secretory traffic, we provided evidence that progression from the early to the late phase of CME is controlled by cargo (Fig. 8A). It is possible that the cell polarity machinery, composed of proteins such as Cdc42 and Cdc24, or other polarized molecules, such as phosphatidylserine, might directly trigger the transition from the early to the late phase of CME (Fairn et al., 2011). However, to date, no study has directly implicated Cdc42 in the molecular mechanism of CME, and phosphatidylserine has been demonstrated to play a role in CME site initiation, but not maturation (Sun and Drubin, 2012). We instead favor the possibility that cargo plays a direct role in CME site maturation, in part because CME would be more efficient with a cargo sensing mechanism. Our studies also suggest that a portion of the variability observed for early arriving endocytic proteins might be explained by differences in the polarization state of the cells analyzed. Thus, in *S. cerevisiae*, cargo regulates CME progression by promoting maturation of stalled endocytic sites rather than by reducing the frequency of abortive sites.

While the specific molecular mechanism of this proposed cargo checkpoint is incompletely understood, several intriguing possibilities provide exciting avenues for further research. Importantly, several studies have suggested that a cargo checkpoint might regulate progress through the mammalian CME pathway (Puthenveedu and von Zastrow, 2006; Mettlen et al., 2010; Loerke et al., 2009; Liu et al., 2010). A recent reconstitution study of mammalian CME suggests that clathrin continually assembles and disassembles during the early stage of the process, with cargo binding stabilizing membrane-associated clathrin structures (Chen et al., 2019). Fluorescently-tagged early arriving CME proteins in budding yeast also fluctuate in intensity, suggesting that such a dynamic proofreading mechanism may be evolutionarily conserved (Carroll et al., 2012). Cargo sensing might involve post-translational modification. Phosphorylation of endocytic proteins by the casein kinase Hrr25 has a documented role in the early stages of CME, and ubiquitination has also been suggested to play a role in CME (Weinberg and Drubin, 2014; Peng et al., 2015). One attractive possibility is that early arriving CME proteins dynamically bind and unbind nascent endocytic sites until cargo binding induces a conformational change, facilitating post-translational modification and maturation.

Because early phase CME proteins do not appear polarized in budding yeast while late phase CME proteins do, we assessed polarization state of more proteins to precisely define which CME proteins are present on either side of the transition point in the pathway. We found that sites containing Ede1, Syp1, Hrr25, the AP-2 complex, Pal1, Sla2, and the epsins all behaved similarly in that they are found both proximal and distal to sites of polarized growth. In contrast, sites containing Pan1 and Sla1, late coat proteins, are highly polarized (Fig. 8A-B). It is interesting to consider implications of our new observations on previous observations. Acute depletion of late coat proteins Pan1 and End3, the first proteins to arrive at CME sites after Sla2 and the epsins, leads to arrest of early endocytic sites on the plasma membrane and aberrant assembly of the late endocytic proteins including the actin cytoskeleton in the cytoplasm (Sun et al., 2015). Thus recruitment of these late coat proteins is likely the crucial step in transitioning from the early phase to the late phase of CME. Conversely, deletion of genes encoding seven early arriving proteins (Syp1, Ede1, Apl1, Pal1, Pal2, Yap1801, and Yap1802) leads to defects in cargo collection but does not halt endocytosis itself, instead shortening the process, consistent with removal of a cargo checkpoint (Brach et al., 2014). This phenotype is reminiscent of the *ede1Δ* phenotype, suggesting that Ede1 may play a direct role in licensing CME site maturation (Stimpson et al., 2009). Loss of a “checkpoint” protein that halts progress in endocytosis in the absence of cargo may be permissive for progress in the CME pathway if that protein does not perform an additional, essential CME function. One of these seven early arriving proteins might therefore be responsible for detecting cargo in a nascent endocytic pit and releasing the block on Pan1/End3 recruitment.

Together, the data we collected and analyzed in this study suggest a conceptual model to explain where and when endocytic internalization is triggered (Fig. 8B). Endocytic sites are initiated widely across the plasma membrane. However, in the absence of cargo delivery, they mature to the internalization phase slowly due to an unsatisfied cargo checkpoint. Polarized delivery of endocytic cargos via exocytosis triggers localized maturation of CME sites through release of the cargo checkpoint. This interpretation represents an attractive explanation for the decades-old observation that “actin patches” (late CME sites) are polarized in small- and medium-budded yeast cells (Adams and Pringle, 1984). Because secretion is directed primarily to the bud while it is growing (Field and Schekman, 1980), concentrated endocytic cargos in the bud trigger local maturation of endocytic sites through the cargo checkpoint, after which actin assembles.

It is interesting to note both similarities and differences between the mechanism of CME regulation described here and the mechanism that has been proposed for mammalian cells. Like the mechanism we describe here, CME sites in mammalian cells are also thought to initiate widely across the membrane but mature to the internalization phase in response to cargo (Ehrlich et al., 2004; Loerke et al., 2009). In contrast to our findings, however, CME sites in mammalian cells are thought by some to be abortive, rather than stalled, in the absence of cargo. The existence of abortive CME sites is controversial, and strictly speaking the mode of regulation described here could also be at play in mammalian cells, even if abortive events are prevalent. Interestingly, activated PDZ-domain interacting G protein-coupled receptors (GPCRs) have been reported to slow CME by a stalling mechanism similar to the one we report here, wherein CME sites containing the activated GPCRs advance more slowly to the internalization phase of the process (Puthenveedu and von Zastrow, 2006). It will be interesting in the future to further compare and contrast mechanisms of cargo-regulated CME between diverse organisms.

## Materials and methods

### Strains and yeast husbandry

The strains used in this study are listed in Table S1. All budding yeast strains were derived from the wild-type diploid DDY1102 and propagated using standard techniques (Amberg et al., 2005). C-terminal fluorescent protein fusions were constructed specifically for this study or for earlier studies, as indicated in Table S1, as described previously (Longtine et al., 1998).

### Live-cell imaging

Cells were grown to mid log phase in imaging media (synthetic minimal media supplemented with Adenine, L-Histidine, L-Leucine, L-Lysine, L-Methionine, Uracil, and 2% glucose), then adhered to coverslips coated with 0.2 mg/ml Concanavalin A.

TIRFM imaging was performed on a Nikon Eclipse Ti2 inverted microscope with a Nikon CFI60 60× 1.49 numerical aperture (NA) Apo oil immersion TIRFM objective and a Hamamatsu Orca-Flash 4.0 V2 sCMOS camera. GFP and RFP/mCherry were excited using 488- and 561-nm lasers and detected using a Chroma HC TIRFM Quad Dichroic (C-FL TIRF Ultra Hi S/N 405/488/561/638) and Chroma HC Quad emission filters BP 525/550 and BP 600/650, respectively. Channels were acquired sequentially. The system was controlled with Nikon Elements software and maintained at 25°C by an OkoLab environmental chamber. Frames were separated by 1 sec.

Epifluorescence imaging was carried out on a Nikon Eclipse Ti inverted microscope with a Nikon 100x 1.4 NA Plan Apo VC oil immersion objective and an Andor Neo 5.5 sCMOS camera. GFP and RFP/mCherry fluorescence were excited using a Lumencore Spectra X LED light source with 550/15 nm and 470/22 nm excitation filters. For two-color imaging, channels were acquired sequentially using an FF-493/574-Di01 dual pass dichroic mirror and FF01-524/628-25 dual pass emission filters (Semrock, Rochester, NY). The system was controlled with Nikon Elements software and maintained at 25**°**C by an environmental chamber (In Vivo Scientific, St. Louis, MO).

Spinning disc confocal microscopy was carried on a Nikon Eclipse Ti inverted microscope with a Nikon 100x 1.45 NA Plan Apo λ oil immersion objective, an Andor IXon X3 EM-CCD camera and Andor CSU-X spinning disc confocal equipment. GFP fluorescence was excited using a 488 nm laser and detected with a Chroma 535/20 nm emission filter (Bellows Falls, VT). mCherry fluorescence was excited using a 565 nm laser and detected with a Chroma 605/52 nm emission filter. The system was controlled with Nikon Elements software. Imaging was conducted at room temperature (∼23**°**C).

### Osmotic shock and BFA experiments

For osmotic shock and BFA experiments, cells were adhered to Concanavalin A coated coverslips for live cell imaging. For osmotic shock experiments, the cells were overlaid with only 250 µL imaging media. Osmotic shock was achieved through rapid addition of 4 mL of sterile water or 4 mL imaging media as a control. For BFA experiments, the cells were overlaid with 500 µL imaging media and BFA treatment was initiated by rapidly adding imaging media supplemented with BFA such that the final concentration was 75 µg/mL (Graham et al., 1993).

### Image and data analysis

Tracking of endocytic sites was performed using the MATLAB package cmeAnalysis (Aguet et al., 2013). The red and green channels for each movie were tracked independently, and the centroid position was determined for each site over time. These data were used as input in custom MATLAB scripts that associated colocalized tracks in the red and green channels while rejecting sites that appeared within 350 nm of one another (Hong et al., 2015). The fluorescence intensity for each spot was calculated as an integrated intensity within a circular region with a diameter of 7 pixels (756 nm), centered at the spot position determined by cmeAnalysis. The background fluorescence was calculated as the average fluorescence intensity of an annulus 2 pixels (216 nm) wide surrounding the circular region used to calculate the spot intensity.

Custom MATLAB scripts were then used to clean our dataset by rejecting tracks that fell into several categories. Sites with a low signal-to-noise (brightness not significantly above background, Hong et al., 2015) were excluded from further analysis. Tracks that began or terminated within 5 frames of the beginning or end of the movie were excluded so that only complete tracks were kept for analysis. Sites whose calculated background-subtracted intensity went below −0.25× the maximum intensity were excluded to prevent abnormally high background or presence of nearby sites from affecting intensity measurements. Finally, outliers in spot lifetime, fluorescence intensity, time to arrival, and time to disappearance (defined as more than 1.5× the interquartile distance from the median value) were also excluded. This was done to exclude erroneous traces in which two distinct tracks had been inadvertently linked, where two sites overlapped and thus were substantially brighter, as well as erroneous putative colocalizations. While these analyses were blind to cell cycle stage, long-lived early sites in polarized mothers represented a small fraction of the final data set owing to the requirement for association of query protein tracks with reference protein tracks and exclusion of incomplete tracks.

GFP maximum fluorescence intensity and the lifetimes and maximum intensities of the red reference proteins were fit to the lifetimes of the GFP-tagged query proteins via simple linear regression. To compare the slopes of the fit across GFP-tagged proteins with differing brightnesses and lifetimes, the data were first normalized by dividing by their respective medians, which does not change the R^2^ of the fit.

Analysis of the influence of cargo on CME site maturation was carried out using Fiji software (National Institutes of Health, Bethesda, MD). For quantification of Ede1-GFP turnover, circumferential kymographs were generated by drawing a line of width 5 pixels around the circumference of each cell. For quantification of Sla1-GFP and Abp1-RFP patch lifetimes, radial kymographs were generated using a custom Fiji macro that generates a kymograph at every radius around a circle in 2-degree increments. The lifetimes of the first 5 kymographs generated were measured to avoid bias. For intensity profile analysis of Bem1-GFP, intensity profiles were normalized to cell length and averaged using a Fiji plugin provided by C. Brownlee (Brownlee and Heald, 2019). Exact numbers of cells and CME sites measured are described in the figure legends.

### Figure preparation

For figure panels and movies, individual cells were cropped, background fluorescence was uniformly subtracted from the image stack, and photo bleaching was fit to a linear decay function and corrected in Fiji. Look up tables used for display are linear. Plots were generated in MATLAB or Prism 8 (GraphPadSoftware, San Diego, CA). In data dense scatter plots, data points were colorized to indicate data density according to the probability of selecting each point at random based on the estimated underlying probability density function for the data. Figures were assembled in Adobe illustrator Creative Cloud 2019.

### Reproducibility of experiments

All experimental results presented were replicated in at least three distinct experiments to ensure reproducibility.

For TRIFM imaging experiments, endocytic sites from each of the dozens of cells in several fields of view were tracked simultaneously. As the results from each dataset were indistinguishable, they were pooled for further analysis. The number of tracked spots for each protein pair is presented in Tables S2-10. Statistical analysis was conducted in MATLAB.

For epifluorescence and confocal fluorescence imaging experiments, multiple cells from each replicate were analyzed. As data from different replicates were indistinguishable, they were pooled for statistical analysis. The specific number of cells analyzed is indicated in each figure legend. Statistical analysis was conducted in Prism 8.

### Data and code availability

The data and code described can be found at the following links: https://github.com/DrubinBarnes/Pedersen_Hassinger_Marchando_Drubin_CME_Manuscript_2019 https://drive.google.com/drive/folders/1xpDnJ58FxRB7wyPBzLhzPKdyzCjpdNd-?usp=sharing

### Online supplemental materials

Table S1 is a list of yeast strains used in this study. Tables S2-4 are the data plotted in Fig. 1 D-F. Video 1 is the movie used to make the montage in Fig. 1B. Fig. S1 shows lifetime and intensity data from experiments discussed in Figs. 1–4. These data are shown in table form in Tables S5-10. Figs. S2-5 are the complete data sets summarized in Figs. 2–4. These data are shown in table form in Tables S11-15. Videos 2-3 are the movie used to make Fig. 5A-B. Fig. S6 complements Figs. 5–6 and shows analysis of late endocytic site lifetimes during different stages of the cell cycle and before and after osmotic shock, confirmation of depolarization of secretory vesicles upon osmotic shock, and analysis of CME site maturation rates when BFA-insensitive cells are treated with BFA. Fig. S7 shows the polarization state of additional early proteins not shown in Fig. 7 and analysis of late endocytic sites in *7Δ* cells.

## Acknowledgements

We thank Yui Iwamoto for critically reading the manuscript. We are grateful to Tony Bretscher for providing the GFP-Sec4 yeast strain, Michelle Lu for providing the Bem1-GFP yeast strain, and Marko Kaksonen for providing the *7Δ* strains. Spinning disc confocal microscopy was conducted at the University of California, Berkeley Cancer Research Laboratory Molecular Imaging center, supported by the Gordon and Betty Moore foundation. We would like to thank H. Aaron and F. Ives for their microscopy training and assistance. The authors declare no competing financial interests. This research was conducted with US Government support, under and awarded by Department of Defense, Air Force Office of Scientific Research, National Defense Science and Engineering Graduate Fellowship 32 CFR 168a (to J.E.H.); National Institutes of Health Grant R35GM118149 (to D.G.D.).

## Author Contributions

R.T.A. Pedersen, J.E. Hassinger, and D.G. Drubin conceived of the experiments. R.T.A. Pedersen and P. Marchando generated the reagents. R.T.A. Pedersen, J.E. Hassinger, and P. Marchando performed the experiments, and analyzed the data. R.T.A. Pedersen, J.E. Hassinger, and D.G. Drubin wrote the manuscript. D.G. Drubin secured funding.

## Abbreviations

AP-2: Adaptor protein 2
BFA: Brefeldin A
CME: Clathrin-mediated endocytosis
CV: Coefficient of variation
FCho: Fer/CIP4 homology domain only
GFP: Green fluorescent protein
RFP: Red fluorescent protein
TIRFM: Total internal reflection fluorescence microscopy
WASP: Wiskott-Aldrich Syndrome protein
WIP: WASP interacting protein

---

**Figure S1:**
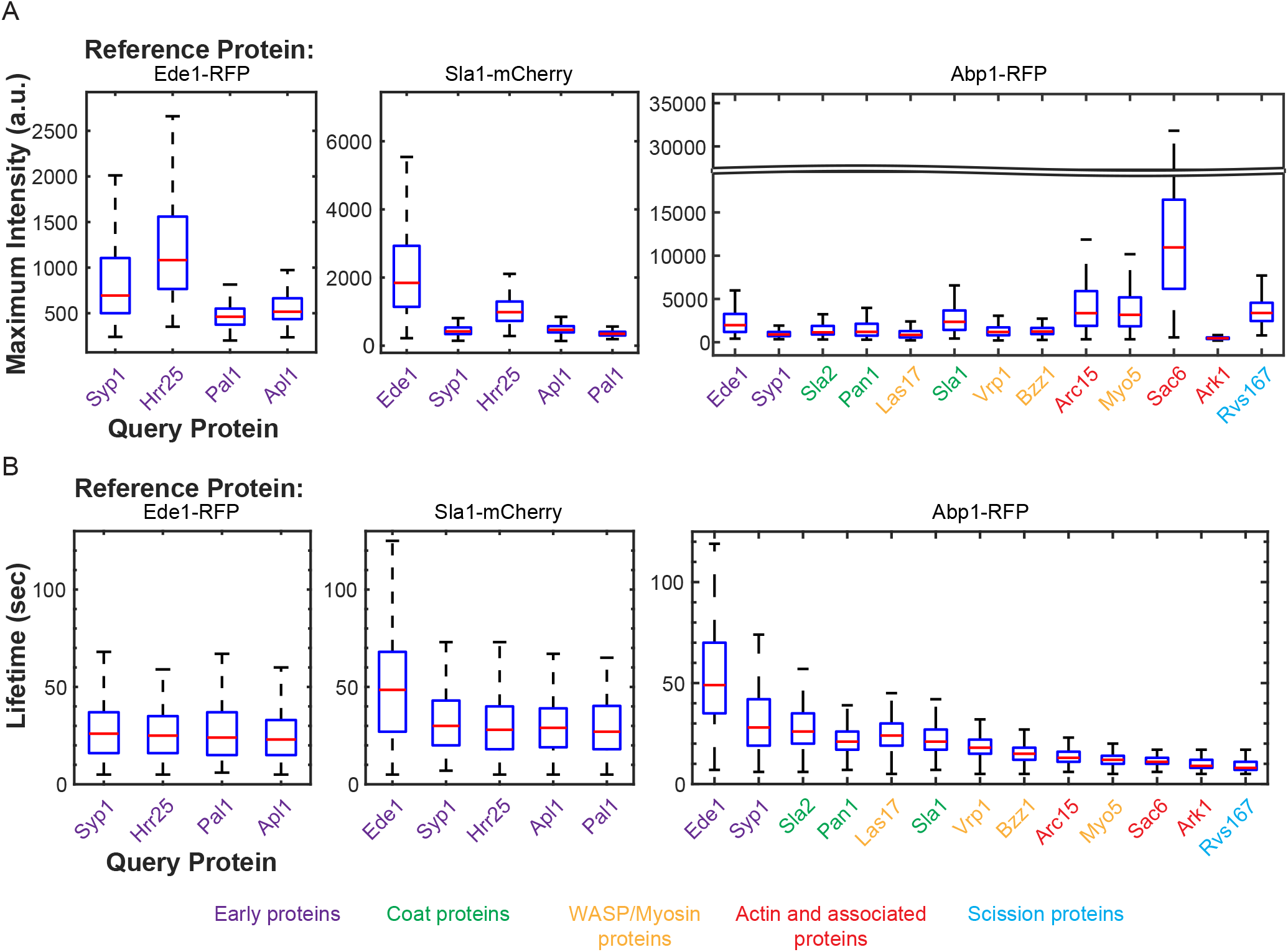
Lifetimes and relative fluorescence intensities of GFP-tagged query proteins agree with prior results but reveal inherent variability. (A) Box and whisker plots of maximum fluorescence intensity of the indicated GFP-tagged query protein imaged with reference to Ede1-RFP (left), Sla1-mCherry (center), and Abp1-RFP (right). (B) Box and whisker plots of lifetimes of the indicated GFP-tagged query protein when colocalized with Ede1-RFP (left), Sla1-mCherry (center), and Abp1-RFP (right). Red lines are median values, boxes indicate interquartile range, and whiskers indicate full range of the data. Colored text indicates which module each protein imaged belongs to according to the color scheme below.

**Figure S2:**
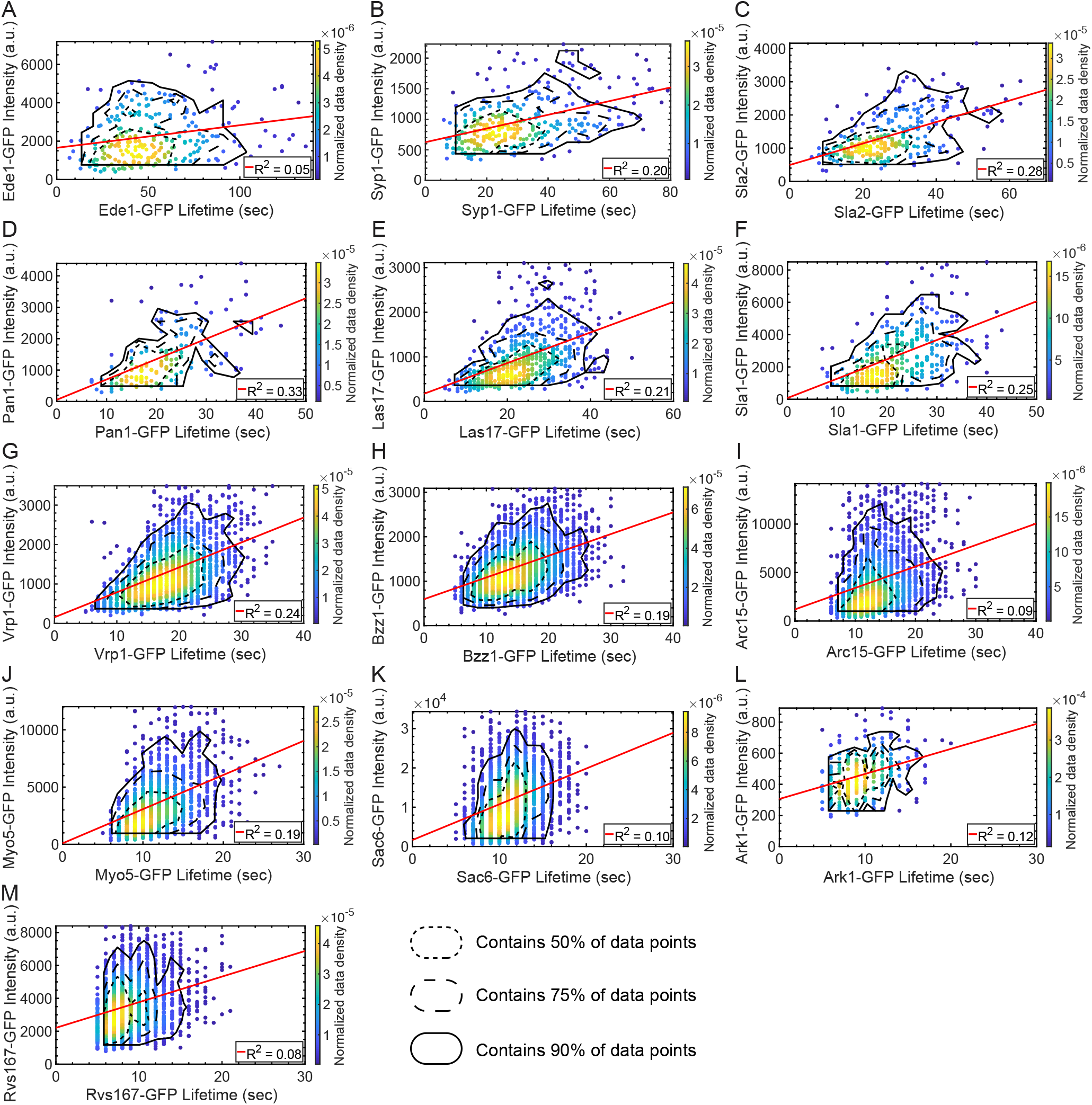
Maximum intensity vs. lifetime plots for GFP-tagged query proteins. Scatter plots of query protein maximum intensity vs. lifetime for GFP-tagged (A) Ede1, (B) Syp1, (C) Sla2, (D) Pan1, (E) Las17, (F) Sla1, (G) Vrp1, (H) Bzz1, (I) Arc15, (J) Myo5, (K) Sac6, (L) Ark1, and (M) Rvs167. Color indicates the normalized data density in the neighborhood around each point. Red lines are linear fits to the data with the indicated R^2^ value. Association of query protein signal with signal of the reference protein Abp1-RFP was used to eliminate spurious events. Contour lines encompassing approximately 50% (short-dashed line), 75% (long-dashed line) and 90% (solid line) encircle local maxima of a ten bin by ten bin histogram of the data.

**Figure S3:**
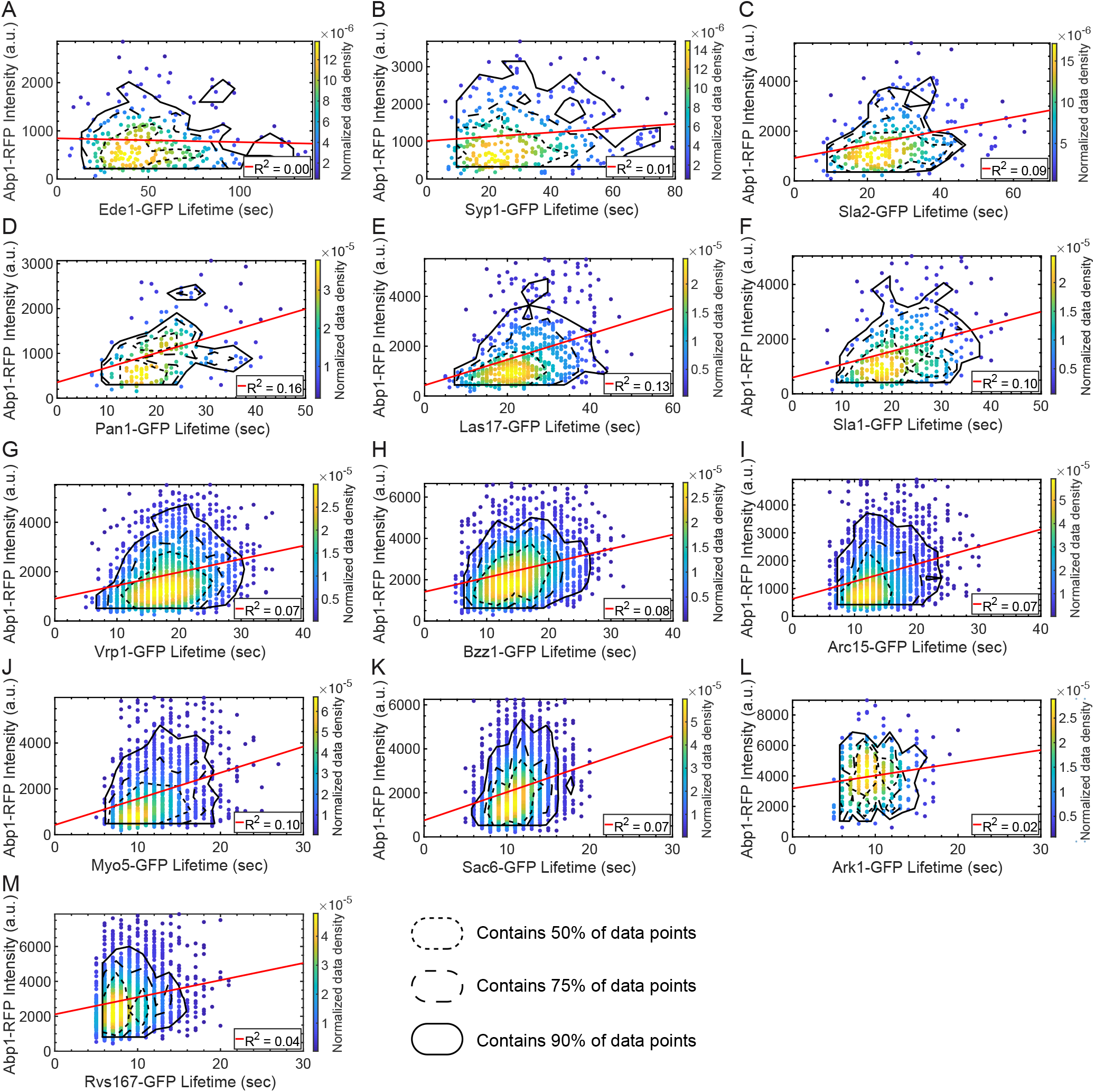
Abp1-RFP maximum intensity vs. query protein lifetime plots for GFP-tagged query proteins. Scatter plots of Abp1-RFP maximum intensity vs. lifetime for GFP-tagged (A) Ede1, (B) Syp1, (C) Sla2, (D) Pan1, (E) Las17, (F) Sla1, (G) Vrp1, (H) Bzz1, (I) Arc15, (J) Myo5, (K) Sac6, (L) Ark1, and (M) Rvs167. Color indicates the normalized data density in the neighborhood around each point. Red lines are linear fits to the data with the indicated R^2^ value. Contour lines encompassing approximately 50% (short-dashed line), 75% (long-dashed line) and 90% (solid line) encircle local maxima of a ten bin by ten bin histogram of the data.

**Figure S4:**
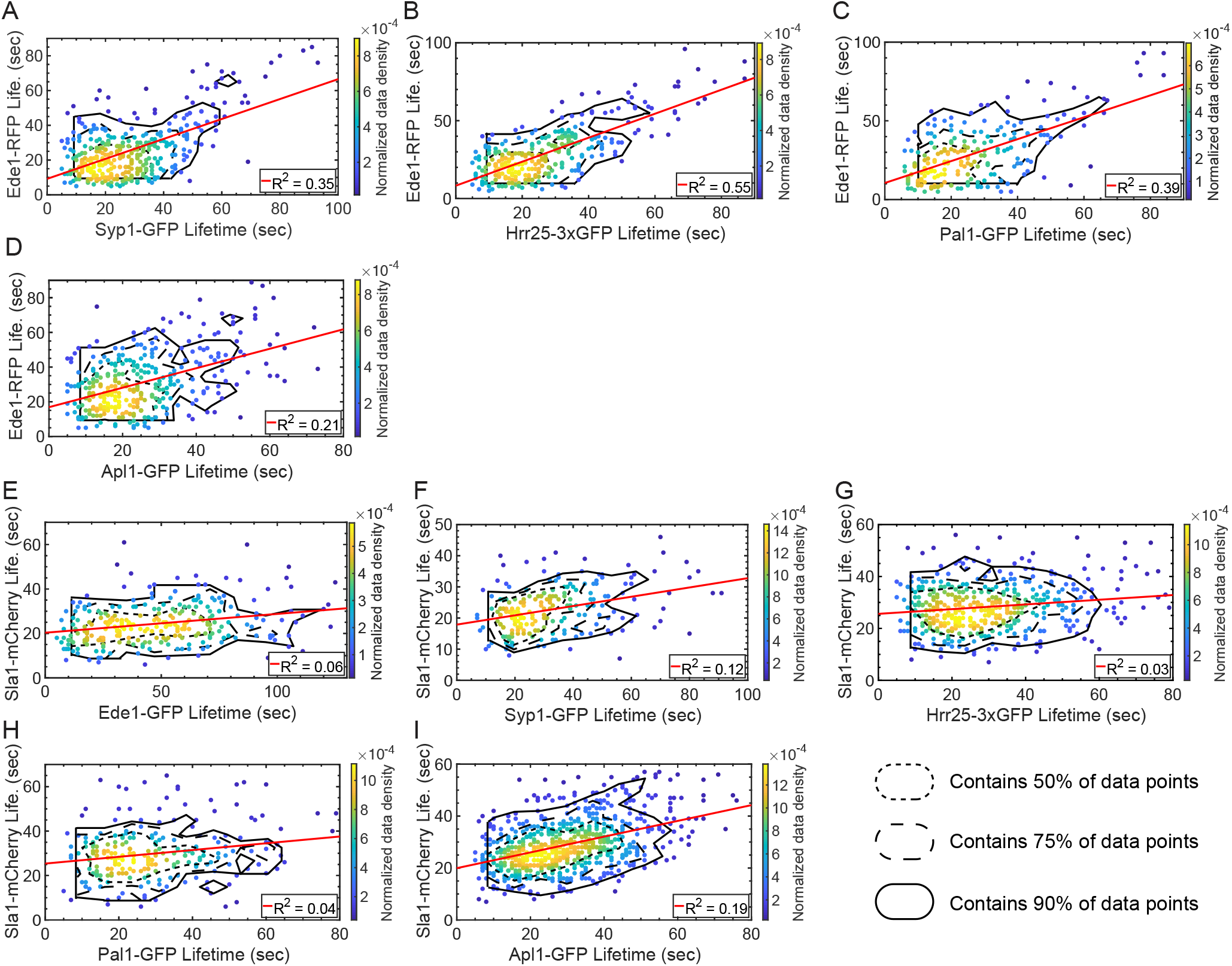
Ede1-RFP lifetime vs. query protein lifetime plots for GFP-tagged query proteins. Scatter plots of Ede1-RFP lifetime vs. query protein lifetime for GFP-tagged (A) Syp1, (B) Hrr25 (3X-GFP tagged), (C) Pal1, and (D) Apl1; and scatter plots of Sla1-mCherry lifetime vs. query protein lifetime for GFP-tagged (E) Ede1, (F) Syp1, (G) Hrr25 (3X-GFP tagged), (H) Pal1, and (I) Apl1. Color indicates the normalized data density in the neighborhood around each point. Red lines are linear fits to the data with the indicated R^2^ value. Contour lines encompassing approximately 50% (short-dashed line), 75% (long-dashed line) and 90% (solid line) encircle local maxima of a ten bin by ten bin histogram of the data.

**Figure S5:**
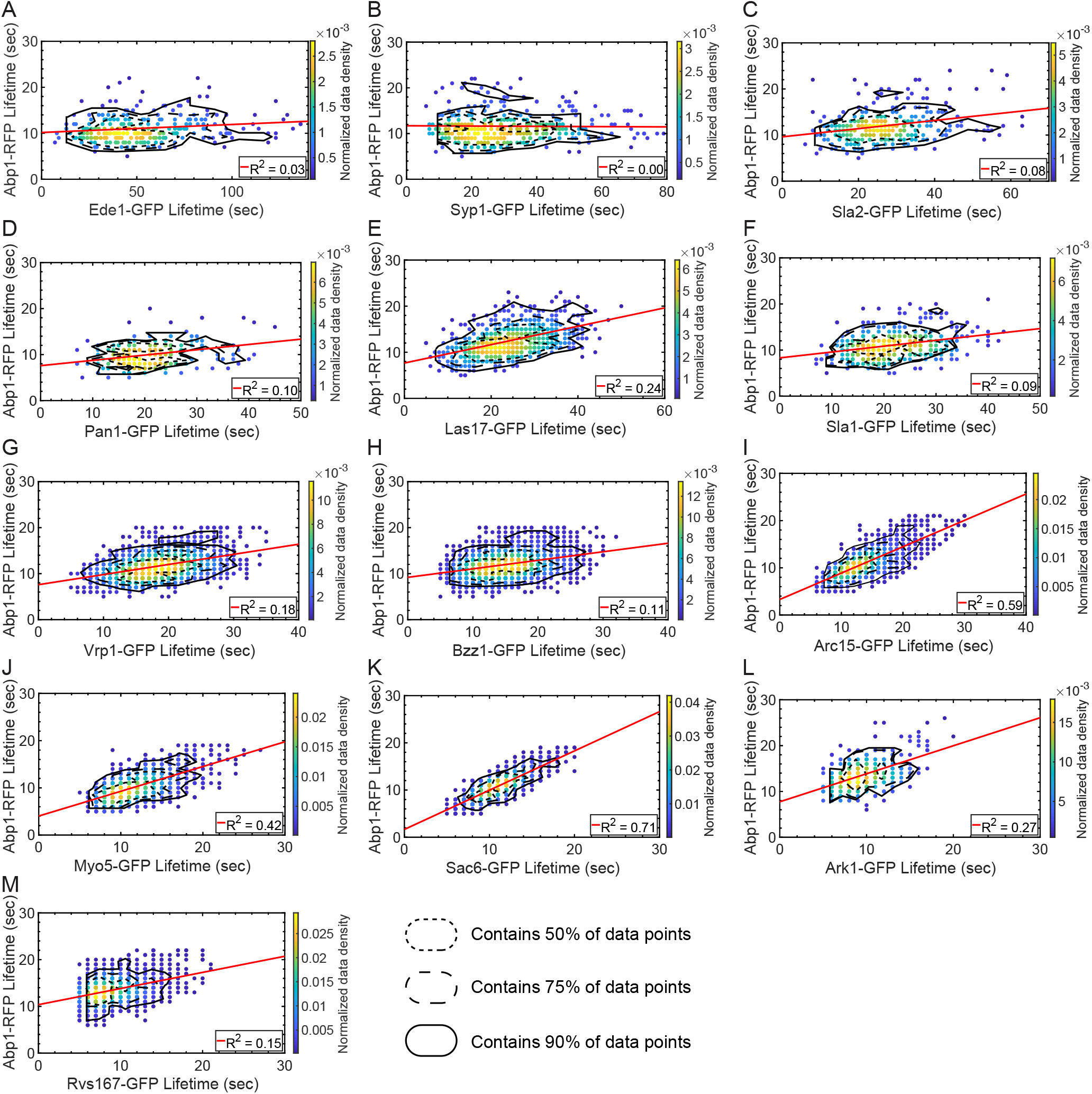
Abp1-RFP lifetime vs. query protein lifetime plots for GFP-tagged query proteins. Scatter plots of Abp1-RFP lifetime vs. query protein lifetime for GFP-tagged (A) Ede1, (B) Syp1, (C) Sla2, (D) Pan1, (E) Las17, (F) Sla1, (G) Vrp1, (H) Bzz1, (I) Arc15, (J) Myo5, (K) Sac6, (L) Ark1, and (M) Rvs167. Color indicates the normalized data density in the neighborhood around each point. Red lines are linear fits to the data with the indicated R^2^ value. Contour lines encompassing approximately 50% (short-dashed line), 75% (long-dashed line) and 90% (solid line) encircle local maxima of a ten bin by ten bin histogram of the data.

**Figure S6:**
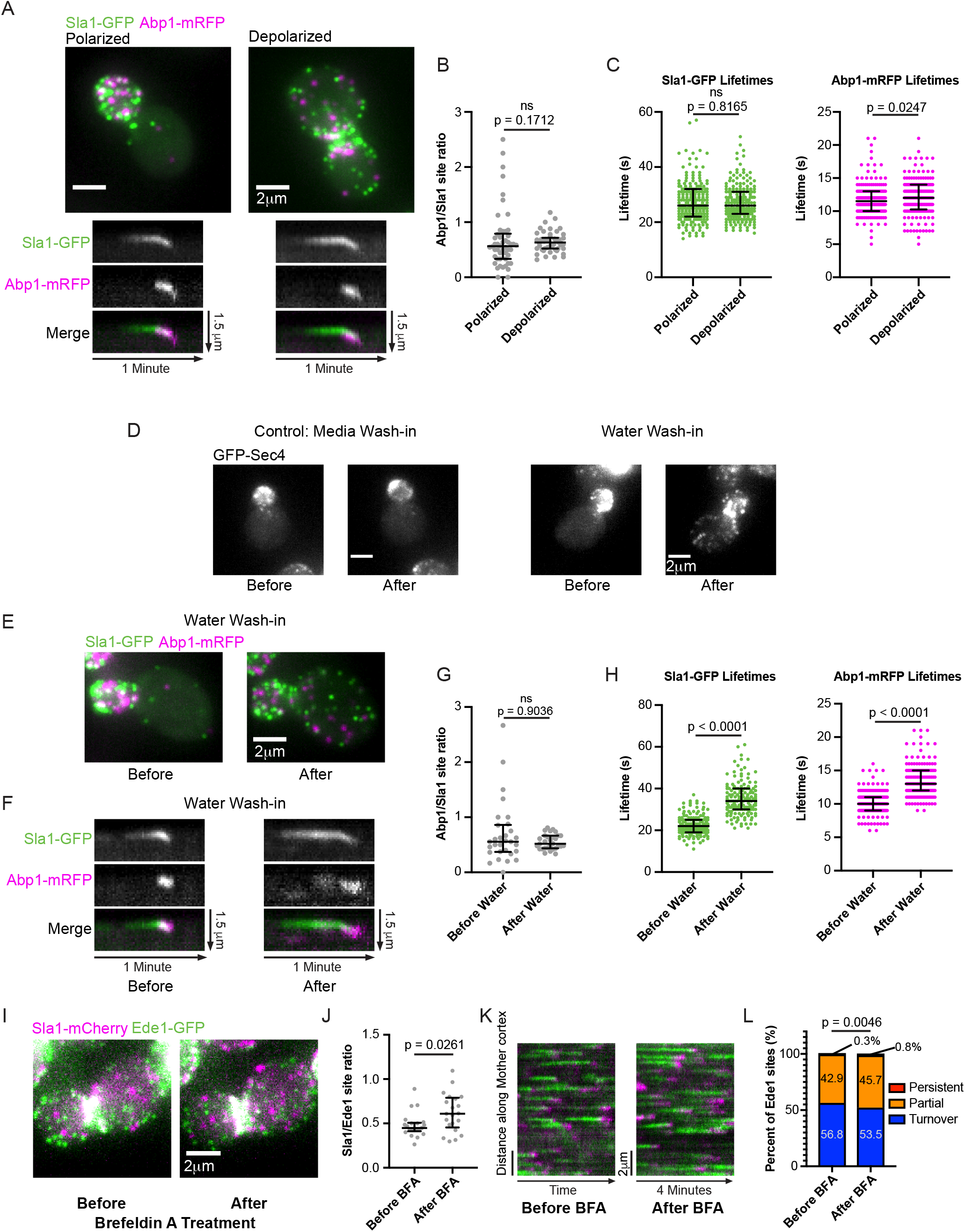
Proximity to sites of exocytosis and cell polarity signaling does not influence the rate of late steps in the endocytic pathway. (A) Maximum intensity projections of z-stacks (top) of polarized and depolarized cells paired with radial kymographs of individual endocytic events from medial focal plane epifluorescence videos of the same cells (bottom) endogenously expressing Sla1-GFP (green) and Abp1-RFP (magenta). Kymograph generation and analysis were carried out in mother cells. (B) Quantification of the ratio of the number of Abp1-RFP sites to Sla1-GFP sites from maximum intensity projections of z-stacks of 50 polarized and 50 depolarized cells. Analysis was carried out in mother cells. A two-tailed p value from a Mann-Whitney U test as in Fig. 5C (U = 1051) is displayed. The median and interquartile ranges are denoted with error bars. (C) Sla1-GFP and Abp1-RFP lifetimes measured in mother cells of 40 polarized cells and 40 depolarized cells (5 representative sites per cell). Two-tailed p values from Mann-Whitney U tests (U_Sla1_ = 19732, U_Abp1_ = 17429) with the null hypothesis that the lifetimes are identical are displayed. The median and interquartile ranges are denoted with error bars. (D) Maximum intensity projections of cells endogenously expressing GFP-Sec4 before and 5 minutes after 17-fold dilution into isotonic imaging media (control, left) or water (right). (E) Maximum intensity projections of a cell endogenously expressing Sla1-GFP (green) and Abp1-RFP (magenta) before and 5 minutes after 17-fold dilution into water. (F) Radial kymographs of individual endocytic events in small-budded mother cells from medial focal plane videos of cells endogenously expressing Sla1-GFP (green) and Abp1-RFP (magenta) before and 5 minutes after 17-fold dilution into water. (G) Quantification of the ratio of the number of Abp1-RFP sites to Sla1-GFP sites from maximum intensity projections of z-stacks of 30 small-budded mother cells before and after osmotic shock. A two-tailed p value from a Mann-Whitney U test as in Fig. 5C (U = 441.5) is displayed. The median and interquartile ranges are denoted with error bars. (H) Sla1-GFP and Abp1-RFP lifetimes measured in 30 small-budded mother cells prior to osmotic shock and 30 cells after osmotic shock (5 representative sites per cell). Two-tailed p values from Mann-Whiney U tests as in (C) (U_Sla1_ = 1678, U_Abp1_ = 2998) are displayed. The median and interquartile ranges are denoted with error bars. (I) Maximum intensity projections of a wild-type cell endogenously expressing Sla1-mCherry (magenta) and Ede1-GFP (green) before and 10 minutes after treatment with Brefeldin A. (J) Quantification of the ratio of the number of Sla1-mCherry sites to Ede1-GFP sites from maximum intensity projections of z-stacks of 21 large-budded mother cells before and after BFA treatment. A two-tailed p value from a Mann-Whitney U test as in Fig. 5C (U = 132.5) is displayed. The median and interquartile ranges are denoted with error bars. (K) Circumferential kymographs around the mother cortex from medial focal plane videos of cells endogenously expressing Sla1-mCherry (magenta) and Ede1-GFP (green) before and 10 minutes after treatment with BFA. (L) Percentage of 375 Ede1 patches from 22 large-budded mother cells before and 477 Ede1 patches from 24 large-budded mother cells after BFA treatment that persist, are partially captured, or turnover during a 4-minute video as in Fig. 5D. A two-tailed p value from a Chi-Square test as in Fig. 5D (chi-square = 10.77, 2 degrees of freedom) is displayed.

**Figure S7:**
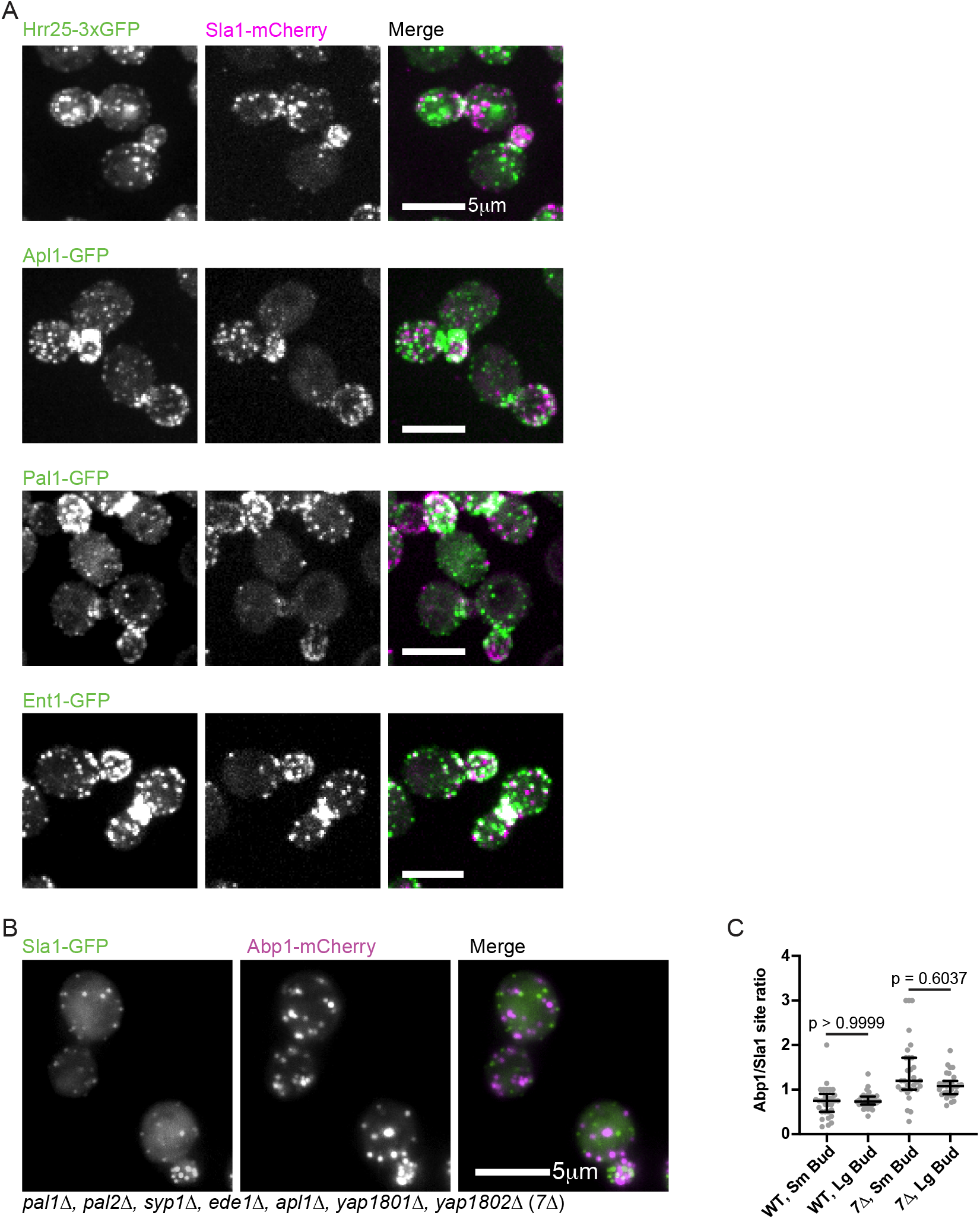
Defining molecular components of the early and late steps of the endocytic pathway. (A) Maximum intensity projections of spinning disc confocal z-stacks of clusters of cells endogenously expressing Sla1-mCherry (magenta) and Hrr25-3xGFP (green, top), Apl1-GFP (green, second row), Pal1-GFP (green, third row), or Ent1-GFP (green, last row). Individual channels are shown in gray scale at left. Images are representative of at least three separate experiments. (B) Maximum intensity projections of epifluorescence z-stacks of clusters of *7Δ* cells endogenously expressing Sla1-GFP (green) and Abp1-mCherry (magenta). Individual channels are shown in gray scale at left. (C) Quantification of the ratio of the number of Abp1-mCherry sites to the number of Sla1-GFP sites in small budded and large budded wild-type and *7Δ* cells (n = 30 cells per category). Numbers are p values from Kruskal-Wallis tests followed by Dunn’s multiple comparisons test. The median and interquartile ranges are denoted with error bars.

**Video 1: Related to Figure 1B**

TIRFM movie of several yeast cells endogenously expressing Las17-GFP (green) and Abp1-RFP (magenta). Each Las17 punctum is punctuated by a burst of Abp1 signal. Frames are separated by 1 second and played back at 15 frames per second (fps).

**Video 2: Related to Figure 5A**

Time-lapse of maximum intensity projections from epifliorescence z-stacks of a cell endogenously expressing Sla1-mCherry (magenta) and Ede1-GFP (green). A dynamic cycle of polarization/depolarization is apparent for the magenta Sla1 sites, but less apparent for the green Ede1 sites. Frames are separated by 10 minutes and played back at 10 fps.

**Video 3: Related to Figure 5B**

Epifluorescence movies in the media focal plane of the polarized and depolarized cells displayed in Figure 5B. The cells endogenously express Sla1-mCherry (magenta) and Ede1-GFP (green). Persistent Ede1 sites are present in the mother of the polarized cell (left), but not in the mother of the depolarized cell (right). Frames are separated by 2 seconds and played back at 15 fps. Circumferential kymographs in Figure 5B were generated from these movies.

**Supplementary table 1:** strains used in this study

| **Name** | **Genotype** | **Source** |
| --- | --- | --- |
| DDY1102 | *MATa/MATα his3-Δ200/his3-Δ200, leu2-3, 112/leu2-3, 112, ura3-52/ura3-52, ade2-1/ADE2, lys2-801/LYS2* | Drubin Lab |
| DDY5713 | *MATa his3-Δ200, leu2-3, 112, ura3-52, ABP1-mRFP::HIS3, SLA1-GFP::KanMX* | This study |
| DDY5714 | *MATa his3-Δ200, leu2-3, 112, ura3-52, ABP1-mRFP::HIS3, Bzz1-GFP::HIS3* | This study |
| DDY5715 | *MATa his3-Δ200, leu2-3, 112, ura3-52, ABP1-mRFP::HIS3, ARC15-GFP::KanMX* | This study |
| DDY5716 | *MATa his3-Δ200, leu2-3, 112, ura3-52, ABP1-mRFP::HIS3, RVS167-GFP::HIS3* | This study |
| DDY5717 | *MATa his3-Δ200, leu2-3, 112, ura3-52, ABP1-mRFP::HIS3, SLA2-GFP::HIS3* | This study |
| DDY5718 | *MATa his3-Δ200, leu2-3, 112, ura3-52, ABP1-mRFP::HIS3, ARK1-GFP::KanMX* | This study |
| DDY5719 | *MATa his3-Δ200, leu2-3, 112, ura3-52, ABP1-mRFP::HIS3, SAC6-GFP::HIS3* | This study |
| DDY5720 | *MATa his3-Δ200, leu2-3, 112, ura3-52, bar1Δ::NatR, EDE1-mRFP::KanMX, PAL1-GFP::HIS3* | This study |
| DDY5721 | *MATa his3-Δ200, leu2-3, 112, ura3-52, bar1Δ::NatR, EDE1-mRFP::HIS3, SYP1-GFP::KanMX* | This study |
| DDY5722 | *MATa his3-Δ200, leu2-3, 112, ura3-52, bar1Δ::NatR, EDE1-mRFP::KanMX, Hrr25-3xGFP::HIS3* | This study |
| DDY5723 | *MATa his3-Δ200, leu2-3, 112, ura3-52, bar1Δ::NatR, SLA1-mCherry::HIS3, PAL1-GFP::HIS3* | This study |
| DDY5724 | *MATa his3-Δ200, leu2-3, 112, ura3-52, bar1Δ::NatR, SLA1-mCherry::HIS3, APL1-GFP::HIS3* | This study |
| DDY5725 | *MATa his3-Δ200, leu2-3, 112, ura3-52, bar1Δ::NatR, SLA1-mCherry::HIS3, EDE1-GFP::HIS3* | This study |
| DDY5726 | *MATa his3-Δ200, leu2-3, 112, ura3-52, bar1Δ::NatR, SLA1-mCherry::HIS3, SYP1-GFP::KanMX* | This study |
| DDY5727 | *MATa his3-Δ200, leu2-3, 112, ura3-52, bar1Δ::NatR, SLA1-mCherry::HIS3, Hrr25-3xGFP::HIS3* | This study |
| DDY3293 | *MATα his3-Δ200, leu2-3, 112, ura3-52, ABP1-mRFP::HIS3, LAS17-GFP::HIS3* | Drubin lab |
| DDY5617 | *MATa his3-Δ200, leu2-3, 112, ura3-52, ABP1-mRFP::HIS3, LAS17-GFP::HIS3* | Drubin lab (Sun et al., 2017) |
| DDY3288 | *MATa his3-Δ200, leu2-3, 112, ura3-52, ABP1-mRFP::HIS3, MYO5-GFP::URA3::KanMX* | Drubin lab |
| DDY5640 | *MATa his3-Δ200, leu2-3, 112, ura3-52, ABP1-mRFP::HygMX, MYO5-GFP::HIS3* | Drubin lab (Sun et al., 2017) |
| DDY3063 | *MATα his3-Δ200, leu2-3, 112, ura3-52, ABP1-mRFP::HIS3, Pan1-GFP::HIS3* | Drubin lab |
| DDY3314 | *MATa his3-Δ200, leu2-3, 112, ura3-52, ABP1-mRFP::HIS3, Pan1-GFP::KanMX* | Drubin lab |
| DDY3867 | *MATα his3-Δ200, leu2-3, 112, ura3-52, ABP1-mRFP::HIS3, SYP1-GFP::KanMX* | Drubin lab (Stimpson et al., 2009) |
| DDY3868 | *MATa his3-Δ200, leu2-3, 112, ura3-52, ABP1-mRFP::HIS3, Ede1-GFP::HIS3* | Drubin lab (Stimpson et al., 2009) |
| DDY3061 | *MATα his3-Δ200, leu2-3, 112, ura3-52, ABP1-mRFP::HIS3, CLC1-GFP::HIS3* | Drubin lab |
| DDY5635 | *MATa his3-Δ200, leu2-3, 112, ura3-52, ABP1-mRFP::HygMX, VRP1-GFP::HIS3* | Drubin lab (Sun et al., 2017) |
| DDY3837 | *MATα, his3-Δ200, leu2-3, 112, ura3-52, bar1Δ::LEU2, EDE1-mRFP::HIS3, APL1-GFP::HIS3* | Drubin lab |
| DDY5728 | *MATα, his3Δ1, leu2Δ0, lys2Δ0, ura3Δ0, GFP-SEC4::URA3* | Bretscher lab (Donovan and Bretscher, 2015) |
| DDY4884 | *MATα, his3-Δ200, leu2-3, 112, ura3-52, BEM1-GFP::HIS3* | Drubin Lab |
| DDY4083 | *MATα, his3-Δ200, leu2-3, 112, ura3-52, SLA1-mCherry::HIS3, SLA2-GFP::HIS3* | Drubin lab (Carroll et al., 2012) |
| DDY4084 | *MATα, his3-Δ200, leu2-3, 112, ura3-52, SLA1-mCherry::HIS3, ENT1-GFP::HIS3* | Drubin lab (Carroll et al., 2012) |
| DDY4085 | *MATα, his3-Δ200, leu2-3, 112, ura3-52, SLA1-mCherry::HIS3, ENT2-GFP::HIS3* | Drubin lab (Carroll et al., 2012) |
| DDY4834 | *MATα, his3-Δ200, leu2-3, 112, ura3-52, SLA1-mCherry::HIS3, PAN1-GFP::HIS3* | Drubin lab (Sun et al., 2015) |
| DDY5734 | *MATα, his3-Δ200, leu2-3, 112, ura3-52, EDE1-GFP::HIS3, SLA1-mCherry::HIS3, hrr25-AID::KanMX, TIR1::LEU2* | This study |
| DDY5735 | *MATα, his3-Δ200, leu2-3, 112, ura3-52, EDE1-GFP::HIS3, SLA1-mCherry::HIS3, hrr25-AID::KanMX* | This study |
| DDY5736 | *MATα, his3-Δ200, leu2-3, 112, ura3-52, EDE1-GFP::HIS3, SLA1-mCherry::HIS3, TIR1::LEU2* | This study |
| DDY5737 | *MATa, his3-Δ200, leu2-3, 112, ura3-52, GEV::URA3, pGAL-HRR25-13Myc::LEU2, EDE1-GFP::HIS3, SLA1-mCherry::HIS3, bar1Δ::NatR* | This study |
| DDY5738 | *MATa, his3-Δ200, leu2-3, 112, ura3-52, pGAL-HRR25-13Myc::LEU2, EDE1-GFP::HIS3, SLA1-mCherry::HIS3, bar1Δ::NatR* | This study |
| DDY5739 | *MATa, his3-Δ200, leu2-3, 112, ura3-52, GEV::URA3, EDE1-GFP::HIS3, SLA1-mCherry::HIS3, bar1Δ::NatR* | This study |
| DDY5730 | *MATa, his3Δ200, leu2-3,112, ura3-52, lys2-801, SLA1-EGFP::HIS3MX6, ABP1-mCherry::kanMX4* | Kaksonen lab (Brach et al. 2014) |
| DDY5731 | *MATa, his3Δ200, leu2-3,112, ura3-52, lys2-801, SLA2-EGFP::natNT2, Sla1-mCherry::kanMX4* | Kaksonen lab (Brach et al. 2014) |
| DDY5732 | *MATa, his3Δ200, leu2-3,112, ura3-52, lys2-801, pal1::natNT2, pal2::natNT2, syp1::hphNT1, ede1::natNT2, apl1::natNT2, yap1801::natNT2, yap1802::natNT2, SLA1-EGFP::HIS3MX6, ABP1-mCherry::kanMX4* | Kaksonen lab (Brach et al. 2014) |
| DDY5733 | *MATa, his3Δ200, leu2-3,112, ura3-52, lys2-801, pal1::natNT2, pal2::natNT2, syp1::hphNT1, ede1::natNT2, apl1::natNT2, yap1801::natNT2, yap1802::natNT2, SLA2-EGFP::HIS3MX6, SLA1-mCherry::kanMX4* | Kaksonen lab (Brach et al. 2014) |

**Supplementary table 2:**
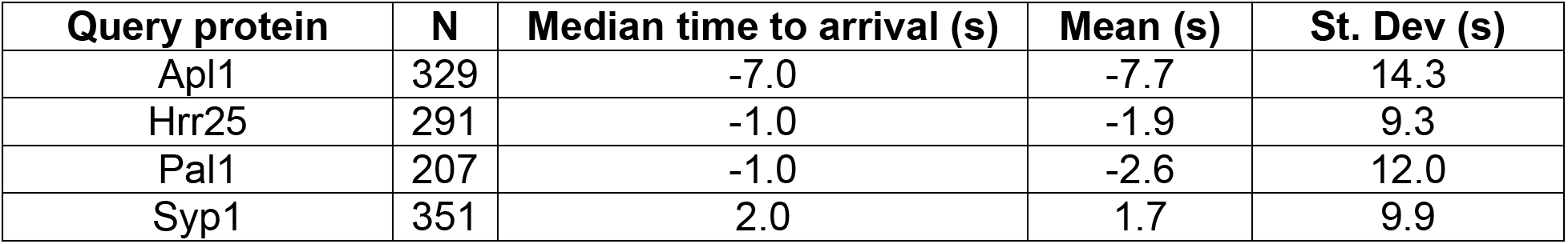
Time to arrival for GFP-tagged query proteins colocalized with Ede1-RFP. Related to Figure 1D

**Supplementary table 3:** Time to arrival for GFP-tagged query proteins colocalized with Sla1-mCherry. Related to Figure 1E

| Query protein | N | Median time to arrival (s) | Mean (s) | St. Dev (s) |
| --- | --- | --- | --- | --- |
| Apl1 | 728 | 3.0 | 3.8 | 12.1 |
| Ede1 | 260 | 29.0 | 29.6 | 26.1 |
| Hrr25 | 444 | 11.5 | 11.3 | 16.5 |
| Pal1 | 229 | 3.0 | 6.0 | 15.7 |
| Syp1 | 211 | 15.0 | 17.4 | 15.3 |

**Supplementary table 4:** Time to arrival for GFP-tagged query proteins colocalized with Abp1-RFP. Related to Figure 1F

| Query protein | N | Median time to arrival (s) | Mean (s) | St. Dev (s) |
| --- | --- | --- | --- | --- |
| Arc15 | 1878 | 2.0 | 2.3 | 2.5 |
| Ark1 | 260 | -2.0 | -2.3 | 2.2 |
| Bzz1 | 2257 | 5.0 | 5.7 | 3.9 |
| Ede1 | 252 | 42.5 | 48.0 | 27.9 |
| Las17 | 582 | 14.0 | 14.1 | 6.9 |
| Myo5 | 1202 | 1.0 | 1.5 | 1.8 |
| Pan1 | 158 | 15.0 | 16.3 | 6.7 |
| Rvs167 | 1802 | -3.0 | -3.1 | 2.0 |
| Sac6 | 2512 | 0.0 | 0.0 | 1.1 |
| Sla1 | 316 | 13.0 | 14.0 | 5.9 |
| Sla2 | 274 | 18.0 | 18.9 | 9.9 |
| Syp1 | 315 | 21.0 | 24.3 | 17.6 |
| Vrp1 | 1604 | 9.0 | 9.0 | 4.4 |

**Supplementary table 5:** Maximum fluorescence intensity of GFP-tagged query proteins colocalized with Ede1-RFP. Related to Figure 1, Figure supplement 1A

| Query protein | N | Median maximum intensity (au) | Mean (au) | St. Dev (au) |
| --- | --- | --- | --- | --- |
| Apl1 | 329 | 517 | 565 | 186 |
| Hrr25 | 291 | 1083 | 1199 | 542 |
| Pal1 | 207 | 462 | 483 | 160 |
| Syp1 | 351 | 694 | 868 | 508 |

**Supplementary table 6:** Maximum fluorescence intensity of GFP-tagged query proteins colocalized with Sla1-mCherry. Related to Figure 1, Figure supplement 1A

| Query protein | N | Median maximum intensity (au) | Mean (au) | St. Dev (au) |
| --- | --- | --- | --- | --- |
| Apl1 | 728 | 466 | 486 | 140 |
| Ede1 | 260 | 1843 | 2183 | 1434 |
| Hrr25 | 444 | 984 | 1052 | 441 |
| Pal1 | 229 | 349 | 363 | 97 |
| Syp1 | 211 | 419 | 461 | 183 |

**Supplementary table 7:** Maximum fluorescence intensity of GFP-tagged query proteins colocalized with Abp1-RFP. Related to Figure 1, Figure supplement 1A

| Query protein | N | Median maximum intensity (au) | Mean (au) | St. Dev (au) |
| --- | --- | --- | --- | --- |
| Arc15 | 1878 | 3358 | 4291 | 3116 |
| Ark1 | 260 | 457 | 465 | 142 |
| Bzz1 | 2257 | 1241 | 1342 | 539 |
| Ede1 | 252 | 1972 | 2292 | 1364 |
| Las17 | 582 | 851 | 1012 | 615 |
| Myo5 | 1202 | 3168 | 3743 | 2457 |
| Pan1 | 158 | 1196 | 1479 | 896 |
| Rvs167 | 1802 | 3378 | 3620 | 1558 |
| Sac6 | 2512 | 10954 | 11916 | 7004 |
| Sla1 | 316 | 2356 | 2749 | 1717 |
| Sla2 | 274 | 1144 | 1399 | 687 |
| Syp1 | 315 | 899 | 979 | 405 |
| Vrp1 | 1604 | 1183 | 1329 | 681 |

**Supplementary table 8:** Lifetimes of GFP-tagged query proteins colocalized with Ede1-RFP. Related to Figure 1, Figure supplement 1B

| Query protein | N | Median lifetime (s) | Mean lifetime (s) | St. Dev (s) | Coeff. of variation |
| --- | --- | --- | --- | --- | --- |
| Apl1 | 329 | 23.0 | 25.7 | 13.9 | 0.54 |
| Hrr25 | 291 | 25.0 | 27.9 | 16.0 | 0.57 |
| Pal1 | 207 | 24.0 | 27.6 | 16.5 | 0.60 |
| Syp1 | 351 | 26.0 | 28.8 | 16.2 | 0.56 |

**Supplementary table 9:** Lifetimes of GFP-tagged query proteins colocalized with Sla1-mCherry. Related to Figure 1, Figure supplement 1B

| Query protein | N | Median lifetime (s) | Mean lifetime (s) | St. Dev (s) | Coeff. of variation |
| --- | --- | --- | --- | --- | --- |
| Apl1 | 728 | 29.0 | 30.0 | 13.7 | 0.46 |
| Ede1 | 260 | 48.5 | 49.5 | 27.4 | 0.55 |
| Hrr25 | 444 | 28.0 | 30.5 | 15.8 | 0.52 |
| Pal1 | 229 | 27.0 | 29.9 | 15.5 | 0.52 |
| Syp1 | 211 | 30.0 | 33.3 | 17.2 | 0.52 |

**Supplementary table 10:**
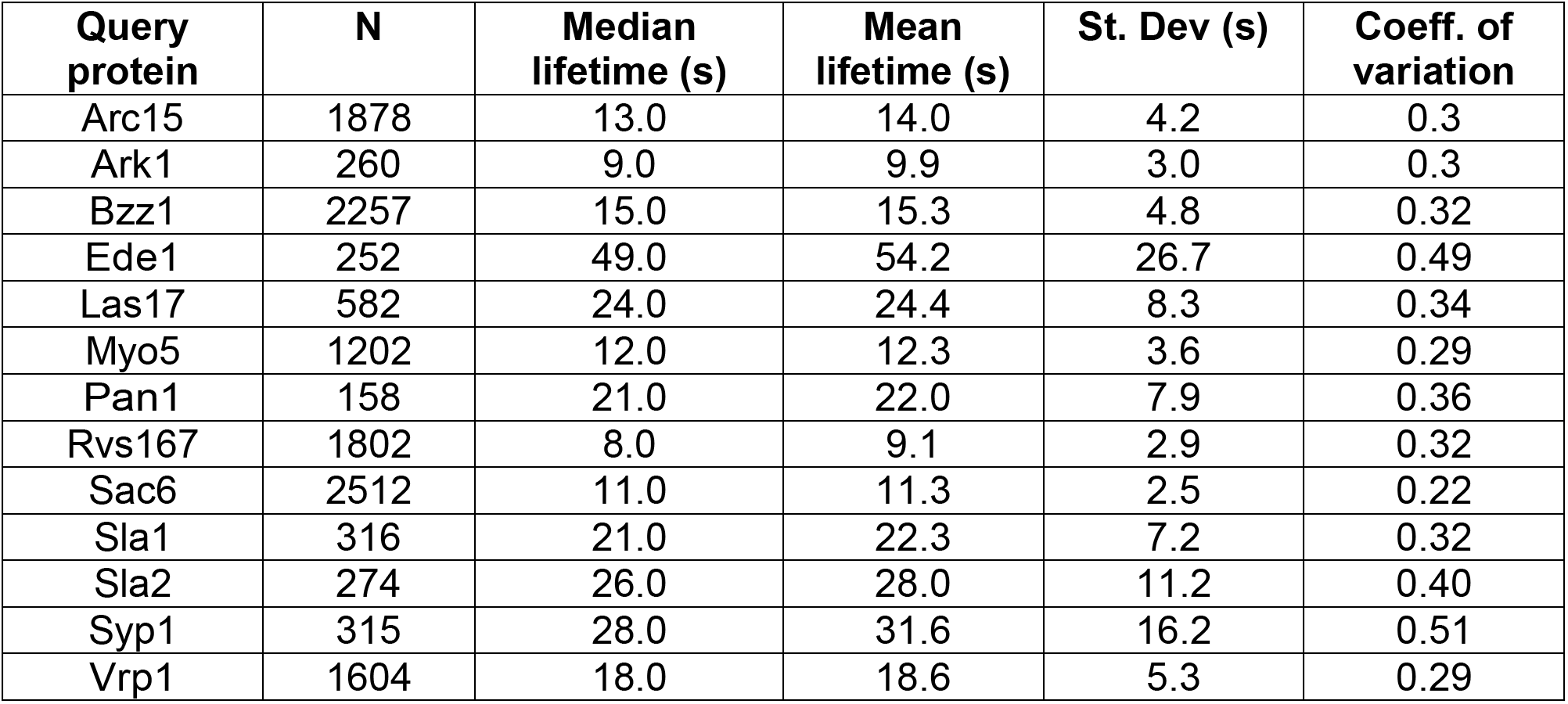
Lifetimes of GFP-tagged query proteins colocalized with Sla1-mCherry. Related to Figure 1, Figure supplement 1B

**Supplementary table 11:** Fit statistics from linear fits of intensity vs. lifetime plots for GFP-tagged query proteins. Upper and lower bounds indicate the 95% confidence interval. Related to Figure 2D

| Query protein | N | Slope | Slope lower bound | Slope upper bound | R<sup>2</sup> | R<sup>2</sup> lower bound | R<sup>2</sup> upper bound |
| --- | --- | --- | --- | --- | --- | --- | --- |
| Arc15 | 1878 | 0.86 | 0.73 | 0.98 | 0.09 | 0.06 | 0.11 |
| Ark1 | 260 | 0.32 | 0.21 | 0.42 | 0.12 | 0.05 | 0.20 |
| Bzz1 | 2257 | 0.59 | 0.54 | 0.64 | 0.19 | 0.16 | 0.22 |
| Ede1 | 252 | 0.29 | 0.14 | 0.45 | 0.05 | 0.01 | 0.12 |
| Las17 | 582 | 0.97 | 0.82 | 1.12 | 0.21 | 0.16 | 0.28 |
| Myo5 | 1202 | 1.13 | 1.00 | 1.26 | 0.19 | 0.15 | 0.23 |
| Pan1 | 158 | 1.14 | 0.88 | 1.39 | 0.33 | 0.21 | 0.45 |
| Rvs167 | 1802 | 0.37 | 0.31 | 0.43 | 0.08 | 0.06 | 0.11 |
| Sac6 | 2512 | 0.91 | 0.81 | 1.02 | 0.10 | 0.08 | 0.13 |
| Sla1 | 316 | 1.07 | 0.86 | 1.27 | 0.25 | 0.17 | 0.33 |
| Sla2 | 274 | 0.73 | 0.59 | 0.88 | 0.28 | 0.19 | 0.37 |
| Syp1 | 315 | 0.35 | 0.27 | 0.43 | 0.20 | 0.13 | 0.28 |
| Vrp1 | 1604 | 0.96 | 0.88 | 1.04 | 0.24 | 0.21 | 0.28 |

**Supplementary table 12:** Fit statistics from linear fits of Abp1-RFP intensity vs. lifetime plots for GFP-tagged query proteins. Upper and lower bounds indicate the 95% confidence interval. Related to Figure 3D

| Query protein | N | Slope | Slope lower bound | Slope upper bound | R<sup>2</sup> | R<sup>2</sup> lower bound | R<sup>2</sup> upper bound |
| --- | --- | --- | --- | --- | --- | --- | --- |
| Arc15 | 1878 | 0.65 | 0.54 | 0.75 | 0.07 | 0.05 | 0.10 |
| Ark1 | 260 | 0.19 | 0.04 | 0.34 | 0.02 | 0.00 | 0.07 |
| Bzz1 | 2257 | 0.46 | 0.39 | 0.52 | 0.08 | 0.06 | 0.10 |
| Ede1 | 252 | -0.06 | -0.24 | 0.12 | 0.00 | 0.03* | 0.01 |
| Las17 | 582 | 0.96 | 0.76 | 1.16 | 0.13 | 0.08 | 0.18 |
| Myo5 | 1202 | 0.95 | 0.79 | 1.11 | 0.10 | 0.07 | 0.14 |
| Pan1 | 158 | 0.73 | 0.47 | 1.00 | 0.16 | 0.07 | 0.27 |
| Rvs167 | 1802 | 0.28 | 0.22 | 0.35 | 0.04 | 0.02 | 0.06 |
| Sac6 | 2512 | 0.70 | 0.60 | 0.80 | 0.07 | 0.05 | 0.09 |
| Sla1 | 316 | 0.74 | 0.49 | 0.98 | 0.10 | 0.05 | 0.17 |
| Sla2 | 274 | 0.50 | 0.31 | 0.70 | 0.09 | 0.03 | 0.16 |
| Syp1 | 315 | 0.16 | 0.01 | 0.30 | 0.01 | 0.00 | 0.05 |
| Vrp1 | 1604 | 0.59 | 0.49 | 0.70 | 0.07 | 0.05 | 0.09 |

\* Since R<sup>2</sup> cannot be negative, the lower bound appears to be above the reported R<sup>2</sup> value. Confidence intervals were calculated on R, which can be negative.

**Supplementary table 13:** Fit statistics from linear fits of Ede1-RFP lifetime vs. lifetime plots for GFP-tagged query proteins. Upper and lower bounds indicate the 95% confidence interval. Related to Figure 4D

| Query protein | N | Slope | Slope lower bound | Slope upper bound | R<sup>2</sup> | R<sup>2</sup> lower bound | R<sup>2</sup> upper bound |
| --- | --- | --- | --- | --- | --- | --- | --- |
| Apl1 | 329 | 0.45 | 0.35 | 0.54 | 0.21 | 0.14 | 0.30 |
| Hrr25 | 291 | 0.71 | 0.64 | 0.79 | 0.55 | 0.47 | 0.62 |
| Pal1 | 207 | 0.64 | 0.53 | 0.75 | 0.39 | 0.29 | 0.49 |
| Syp1 | 351 | 0.65 | 0.55 | 0.74 | 0.35 | 0.27 | 0.43 |

**Supplementary table 14:**
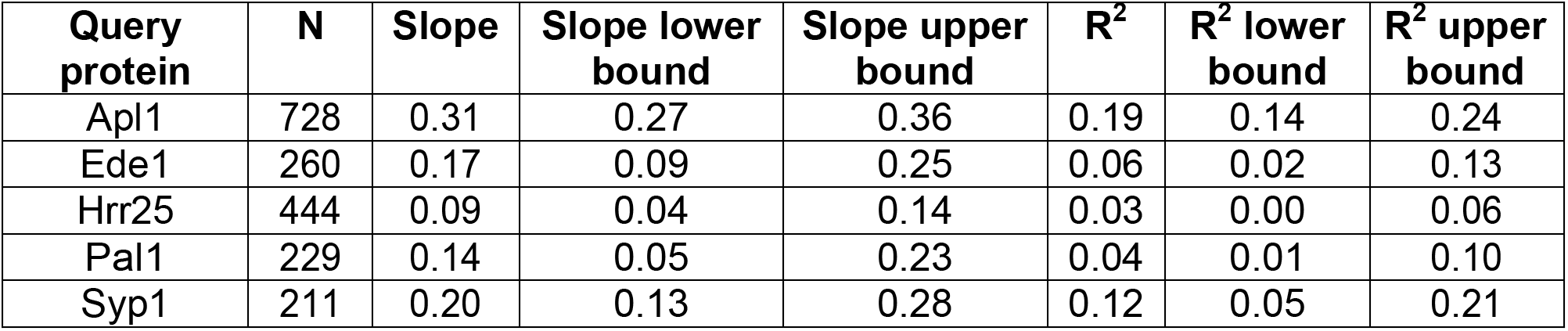
Fit statistics from linear fits of Sla1-mCherry lifetime vs. lifetime plots for GFP-tagged query proteins. Upper and lower bounds indicate the 95% confidence interval. Related to Figure 4E

| Query protein | N | Slope | Slope lower bound | Slope upper bound | R<sup>2</sup> | R<sup>2</sup> lower bound | R<sup>2</sup> upper bound |
| --- | --- | --- | --- | --- | --- | --- | --- |
| Apl1 | 728 | 0.31 | 0.27 | 0.36 | 0.19 | 0.14 | 0.24 |
| Ede1 | 260 | 0.17 | 0.09 | 0.25 | 0.06 | 0.02 | 0.13 |
| Hrr25 | 444 | 0.09 | 0.04 | 0.14 | 0.03 | 0.00 | 0.06 |
| Pal1 | 229 | 0.14 | 0.05 | 0.23 | 0.04 | 0.01 | 0.10 |
| Syp1 | 211 | 0.20 | 0.13 | 0.28 | 0.12 | 0.05 | 0.21 |

**Supplementary table 15:** Fit statistics from linear fits of Abp1-RFP lifetime vs. lifetime plots for GFP-tagged query proteins. Upper and lower bounds indicate the 95% confidence interval. Related to Figure 4F

| Query protein | N | Slope | Slope lower bound | Slope upper bound | R<sup>2</sup> | R<sup>2</sup> lower bound | R<sup>2</sup> upper bound |
| --- | --- | --- | --- | --- | --- | --- | --- |
| Arc15 | 1878 | 0.66 | 0.63 | 0.69 | 0.59 | 0.56 | 0.61 |
| Ark1 | 260 | 0.42 | 0.34 | 0.51 | 0.27 | 0.18 | 0.36 |
| Bzz1 | 2257 | 0.23 | 0.20 | 0.26 | 0.11 | 0.09 | 0.14 |
| Ede1 | 252 | 0.08 | 0.02 | 0.14 | 0.03 | 0.00 | 0.08 |
| Las17 | 582 | 0.40 | 0.34 | 0.46 | 0.24 | 0.19 | 0.31 |
| Myo5 | 1202 | 0.63 | 0.59 | 0.67 | 0.42 | 0.38 | 0.46 |
| Pan1 | 158 | 0.24 | 0.13 | 0.35 | 0.10 | 0.03 | 0.21 |
| Rvs167 | 1802 | 0.21 | 0.19 | 0.24 | 0.15 | 0.12 | 0.18 |
| Sac6 | 2512 | 0.83 | 0.81 | 0.85 | 0.71 | 0.69 | 0.73 |
| Sla1 | 316 | 0.24 | 0.16 | 0.33 | 0.09 | 0.04 | 0.16 |
| Sla2 | 274 | 0.19 | 0.11 | 0.27 | 0.08 | 0.03 | 0.15 |
| Syp1 | 315 | -0.01 | -0.07 | 0.05 | 0.00 | 0.02* | 0.01 |
| Vrp1 | 1604 | 0.36 | 0.32 | 0.40 | 0.18 | 0.14 | 0.21 |

\* Since R<sup>2</sup> cannot be negative, the lower bound appears to be above the reported R<sup>2</sup> value. Confidence intervals were calculated on R, which can be negative.

## Notes

https://github.com/DrubinBarnes/Pedersen_Hassinger_Marchando_Drubin_CME_Manuscript_2019

https://drive.google.com/drive/folders/1xpDnJ58FxRB7wyPBzLhzPKdyzCjpdNd-?usp=sharing

### Competing Interest Statement

The authors have declared no competing interest.

### Summary of Updates

This manuscript replaces the previous revision, uploaded one day earlier, which shows some evidence of having been corrupted during conversion to a PDF.

